# BEAM: a combinatorial recombinase toolbox for binary gene expression and mosaic analysis

**DOI:** 10.1101/2023.02.16.528875

**Authors:** Luciano C. Greig, Mollie B. Woodworth, Alexandros Poulopoulos, Stephanie Lim, Jeffrey D. Macklis

## Abstract

Genetic mosaic analysis, in which mutant cells reside intermingled with wild-type cells, is a powerful experimental approach, but has not been widely used in mice because existing genome-based strategies require complicated and protracted breeding schemes. We have developed an alternative approach termed BEAM (for Binary Expression Aleatory Mosaic) that relies on sparse recombinase activation to generate two genetically distinct, non-overlapping populations of cells for comparative analysis. Following delivery of DNA constructs by transfection or viral transduction, combinatorial recombinase activity generates two distinct populations of cells labeled with either green or red fluorescent protein. Any gene of interest can be mis-expressed or deleted in one population for comparison with intermingled control cells. We have extensively optimized and characterized this system both *in vitro* and *in vivo*, and demonstrate its power for investigating cell autonomy, identifying temporally or spatially aberrant phenotypes, revealing changes in cell proliferation or death, and controlling for procedural variability.

## Introduction

Genetic mosaic analysis of individual mutant cells in an otherwise wild-type organism and local environment offers a powerful approach for investigating gene function with cellular resolution. It can circumvent embryonic lethality, as well as avoiding the confounding pleiotropic effects that often arise in the setting of global loss of gene function throughout an entire organism, or even with conditional loss of gene function in a specific organ, tissue, or cell type(1). By combining genetic mosaic analysis with distinct fluorescent labeling of mutant cells, direct identification of a broad range of abnormal phenotypes becomes possible. Phenotypic analysis can be facilitated further by labeling a second population of wild-type cells with a different fluorescent protein, enabling direct comparison of wild-type and mutant cells within the same tissue.

A number of genome-based strategies for mosaic analysis in mice have been developed. Mosaic analysis with double markers (MADM) relies on interchromosomal recombination of two chimeric alleles, each with a *loxP*-containing intron interrupting the coding sequence for reciprocal halves of GFP and RFP(2). When a null allele for a gene of interest is placed on the same chromosome as the *MADM* allele, CRE-mediated interchromosomal recombination generates green cells that are null for the gene of interest, red cells that are wild-type, and yellow cells that are heterozygous. A second strategy, mosaic analysis with spatial and temporal regulation (MASTR), relies on FLPERT2-mediated excision of an *frt*-flanked transcriptional STOP cassette to activate expression of a GFP-CRE fusion protein(3). Therefore, tamoxifen induction leads to sparse labeling of cells with GFP, as well as efficient recombination by CRE of any *loxP*-flanked (*floxed*) alleles present. These methods have been used in recent years to make important discoveries across a broad range of topics(4)(5)(6)(7)(8). However, they each require time-consuming genetic crosses to enable experimentation, and they each entail additional complexities and limitations. In contrast to genome-based approaches, genetic analysis by delivery of exogenous DNA to defined populations of cells *in vivo* enables versatile manipulation of gene expression, with substantial potential to accelerate biological research in vertebrate model systems. Transfection by chemical or physical means, or viral transduction, can be used to target almost every known cell type in developing or adult animals. In addition, temporal and spatial control over experimental manipulations is implicit, because DNA can be introduced in specific locations and at specific stages of development, maturity, or disease progression. An ever-expanding repertoire of effectors can direct gene knock-down or overexpression, genome editing, cell ablation, modulation of membrane potential, or neuronal circuit tracing, among other possibilities. One often limiting drawback of such approaches relative to genome-based methods, however, is the potential for considerable procedural variability inherent to introduction of exogenous DNA. This variability can cause substantial differences in the number, types, and spatial distribution of cells targeted, often increasing the difficulty of obtaining well-matched pairs of experimental and control specimens, and therefore complicating data analysis and interpretation.

One potential approach to overcoming these limitations is to incorporate an internal control, which would extend the range of feasible experiments and improve the reliability of resulting data. One such approach has been applied to study graded EPHA7/EPHRIN-A signaling in corticothalamic axon sorting within the ventrobasal thalamic nucleus (9). By electroporating a mix of high concentration red fluorescent protein (*RFP*) expression plasmid and low concentration green fluorescent protein (*GFP*)/*EphA7* bicistronic expression plasmid, the authors compared spatial distribution of RFP-positive and RFP/GFP-double-positive axons within the same brain, eliminating the variability introduced by differences in the areal distribution of electroporated neurons across surgeries. Another such approach is to mix a high concentration of *RFP* expression plasmid and of a CRE-dependent *GFP* expression plasmid, together with a low concentration of a *Cre* expression plasmid, resulting in a subset of cells expressing *GFP*(10). Both approaches result in a population of control cells that is labeled with RFP, and a subpopulation of experimental cells that is double-labeled with both RFP and GFP, but neither approach yields truly binary gene expression. Reporter constructs have been previously described that convert from *RFP* expression to *GFP* expression after CRE-mediated recombination(11)(12). However, initial expression of *RFP* in all cells, and/or incomplete recombination, result in a large number of cells co-expressing both fluorescent proteins(13)(14)(15)(16), thus producing potentially ambiguous results.

Here, we substantially extend the range of tools available for genetic mosaic analysis in mice by developing a modular recombinase-based system for binary gene expression and mosaic analysis that can be delivered by transfection or transduction directly into wild-type or floxed mice, without need for complex breeding schemes. The system relies on combinatorial recombinase activation to generate two genetically distinct fluorescently-labeled populations of cells for comparative analysis. Any gene of interest can be misexpressed or deleted in GFP-positive cells, and compared with interspersed RFP-positive control cells. We present its application on previously published loss- and gain-of-function phenotypes to illustrate how BEAM can be employed to great advantage to investigate cell autonomy, to identify temporally or spatially aberrant phenotypes, to reveal changes in cell proliferation or death, and to control for procedural variability. We also applied BEAM to investigate a long-standing hypothesis in the field of cortical development– that each neural progenitor and its progeny constitutes an independent radial unit, and that subsequent organization of the cerebral cortex into functional areas is determined in these neural progenitors as a cell identity protomap that is then transferred to their progeny within a cortical column.

## Results

### Combinatorial recombinase activity can be used to generate genetically distinct populations of cells

Delivery of DNA by transfection, electroporation, and viral transduction is a stochastic process in which each cell can receive anywhere from 0 to >100 copies of the delivered DNA construct, depending on the plasmid concentration or viral titer used, with an approximately normal distribution of copy numbers across the entire population of cells(17). It is, therefore, not possible to use incompatible recombinase sites to generate fully distinct outcomes on a cell-by-cell basis(18), since each possible outcome of recombination is represented multiple times within most cells. However, the proportion of cells that is successfully transfected can be controlled by adjusting plasmid concentration or viral titer. When one plasmid or virus is introduced at higher concentration and another at lower concentration, most cells will receive both or only the one at higher concentration (while a much smaller number of cells will receive only the one at lower). We have exploited this principle, in conjunction with the approximately all-or-none activity of recombinases, to generate binary gene expression outcomes in individual cells (Figure 1A).

**Figure 1.**
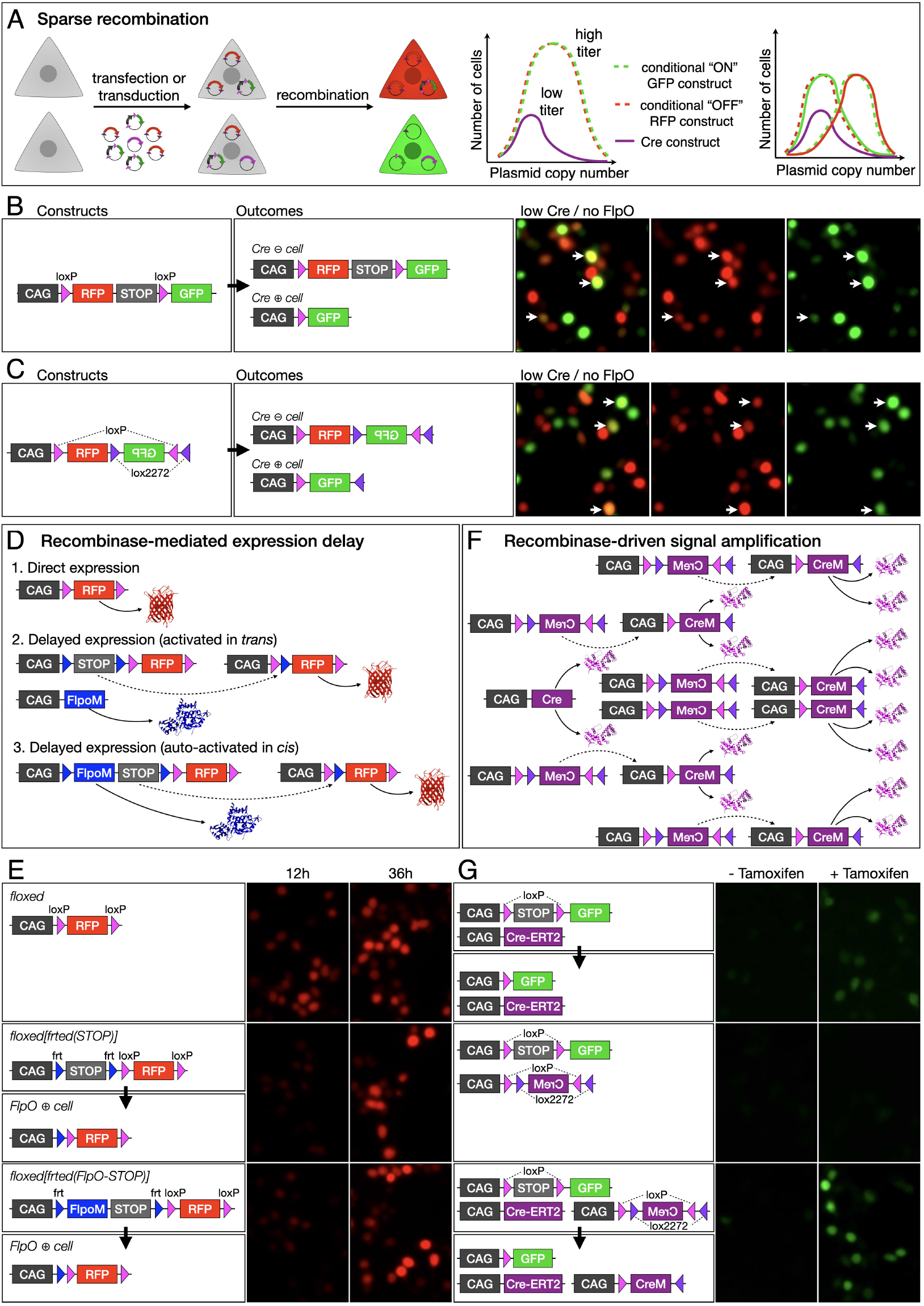
A recombinase-based strategy for binary gene expression. (A) Sparse recombination of conditionally activated DNA constructs results in binary expression outcomes according to construct copy number. Two plasmids, *RFP* activated by CRE and *GFP* inactivated by CRE, are introduced to cells at high titer, together with a *Cre* plasmid at low titer (left). Those cells that receive *Cre* fluoresce green (GFP-on, RFP-off), while those that do not receive *Cre* fluoresce red (GFP-off, RFP-on) (right). (B-C) Transfection of 293T cells with low *Cre* and conditional *RFP*-*GFP* expression constructs results in overlapping expression of *GFP* and *RFP*. Significant overlap of RFP and GFP (arrows, right) result whether using a *loxP-RFP-STOP-loxP-GFP* construct (B) or a *RFP-FLEX(GFP)* construct (C). (D-E) Delayed fluorophore expression. *RFP* can be unconditionally expressed from a *CAG* promoter (B1), or its expression can be delayed by requiring excision of a *STOP* cassette flanked by *frt* sites as a result of *FlpO* expression from a second construct, in *trans* (B2). The delayed expression can be auto-activated in *cis* by flanking both the coding sequence for *FlpO* recombinase and the *STOP* cassette with *frt* sites, thus causing FLPO to excise itself and activate *RFP* expression in one step (B3). Constitutively active *loxP*-flanked *RFP* is expressed at 12 hours post-transfection in 293T cells, with strong expression at 36 hours (E1). Introducing *frt* sites flanking a *STOP* cassette, then co-transfecting with *CAG-FlpO*, reduces *RFP* expression levels at 12 hours post-transfection (E2). Similarly, a self-excising cassette consisting of the coding sequence for *FlpO* and a transcriptional *STOP* also results in delayed expression of *RFP* (E3). (F-G) Recombinase signal amplification. *Cre* expression can be amplified in *trans* by transfecting a low-dose *CAG-Cre* that activates multiple copies of a *FLEX* construct, in which the coding sequence of an intron-containing *Cre* (*CreM*) is reversed. We used a low dose of tamoxifen to induce *GFP* expression in 293T cells transfected with a *Cre-ERT2* expression construct and a *loxP-STOP-loxP-GFP* reporter, obtaining low levels of GFP (G1). Transfection with a *FLEX(CreM)* construct did not result in significant auto-activation in the presence or absence of tamoxifen (G2). However, in the presence of CRE-ERT2 and tamoxifen, *FLEX(CreM)* amplified the recombination signal, substantially increasing GFP levels.

The simplest version of a reporter construct for this purpose would be comprised of a *CMV/*β*-actin* hybrid promoter (*CAG*) and a *loxP*-flanked (*floxed*) *RFP* and transcriptional *STOP* cassette, followed by *GFP*(11)(13). At baseline, the construct drives expression of *RFP*, but *GFP* is expressed after Cre deletes *RFP* (Figure 1B). Alternatively, the same outcome can be achieved using a *CAG*-driven FLip EXcision inverted *lox* site (FLEX) construct, in which the coding sequence for *RFP* is oriented in the sense direction, and the coding sequence for *GFP* is oriented in the antisense direction(12)(14). In this case, CRE permanently reverses the orientation of *GFP* and, in the process, excises the coding sequence for *RFP* (Figure 1C). By co-transfecting each of these constructs with a low dose of *CAG-Cre*, it is possible to generate cells that are primarily green or primarily red. However, there is still substantial overlap in expression, which results from two main limitations: 1) these constructs drive expression of RFP until CRE mediates recombination; and 2) low *Cre* expression results in incomplete recombination of the multiple copies of reporter plasmid in each cell.

To address the first limitation of these prior methods, we sought to delay initial expression of *RFP*. We designed a construct with a *CAG* promoter and an *frt*-flanked (*frted*) transcriptional *STOP* cassette followed by a *floxed*(*RFP*) (Figure 1D), resulting in no expression of *RFP* at baseline, until it is activated by FLP-mediated excision of the *frted*-*STOP* cassette. When cells are transfected with this construct, together with a *CAG-FlpO* construct, *RFP* is expressed, but with a delay introduced by the requirement for *FlpO* transcription and translation, followed by excision of the *frted*(*STOP*) cassette. We also generated a construct with an intron-modified *FlpO* (*FlpoM*) between the first *frt* site and the *STOP* cassette (Figure 1D). *FlpoM* contains a *beta-globin* intron that interrupts its coding sequence and prevents FLPO production in bacteria, thereby avoiding recombination of *frt* sites in *E. coli* as the plasmid is being produced. Once introduced into eukaryotic cells, the plasmid drives transcription of *FlpoM*, the *beta-globin* intron is spliced out, FLPO protein is produced and excises its own coding sequence, along with the transcriptional *STOP* cassette from the plasmid, enabling expression of *RFP*. Therefore, the plasmid functions as a self-activating recombinase-operated delay switch.

In 293T cells transfected with *FlpO*-delayed expression constructs, RFP fluorescence is clearly reduced 12 hours after transfection, but approaches similar levels to cells transfected with a direct expression construct by 36 hours after transfection (Figure 1E). When these plasmids are co-transfected with a *CAG-Cre* construct, there is direct competition between CRE-mediated excision of *RFP*, and FLPO-mediated excision of the *STOP* cassette, leading to reduced RFP fluorescence in CRE-positive cells (Figure S1A). Therefore, this approach substantially mitigates one of the mechanisms that lead to overlap between GFP and RFP in a *Cre*-dependent binary expression system.

To address the second limitation of existing methods, that of incomplete recombination, we developed a strategy for recombinase-mediated signal amplification. We cloned an intron-modified *Cre* (*CreM*) into a *CAG*-driven *FLEX* cassette in the antisense orientation, thereby generating a CRE-activated *Cre* expression construct in which the enzymatic activity of CRE recursively activates expression of more *Cre* in a self-amplifying reaction (Figure 1F). We tested this approach by transfecting 293T cells with a *Cre-ERT2* expression construct and a *floxed(STOP)-GFP* reporter, then inducing recombination with a low concentration of tamoxifen (Figure 1G). This results in recombination of only a subset of the available copies of *floxed(STOP)-GFP* construct in each cell, and therefore low levels of GFP fluorescence. Co-transfecting the *FLEX(CreM)* construct along with the *Cre-ERT2* construct results in higher levels of GFP fluorescence. Importantly, we do not observe leaky auto-activation of *FLEX(CreM)* in the absence of *Cre-ERT2*, indicating that it remains inert in mammalian cells in the absence of initiating CRE activity from a second source.

### BEAM drives binary gene expression *in vitro* and *in vivo*

Combining both of these advances– *FlpO*-delayed *RFP* expression and amplification of CRE activity with *FLEX(CreM)*– dramatically reduced overlap between EGFP and RFP (Figure 2A). We used FACS to quantify the extent to which each of these approaches can sharpen binary expression. By introducing *FlpO*-delayed *RFP* expression, we find a substantial reduction in the overlap of EGFP and RFP from 20.3% to 8% (Figure 1H). Strikingly, by combining *FlpO*-delayed *RFP* expression with CRE signal amplification, overlap between EGFP and RFP is reduced to only 3.6%. This combination of constructs forms the basis of our Binary Expression Aleatory Mosaic (BEAM) recombinase system, and are used in subsequent experiments unless otherwise noted.

**Figure 2.**
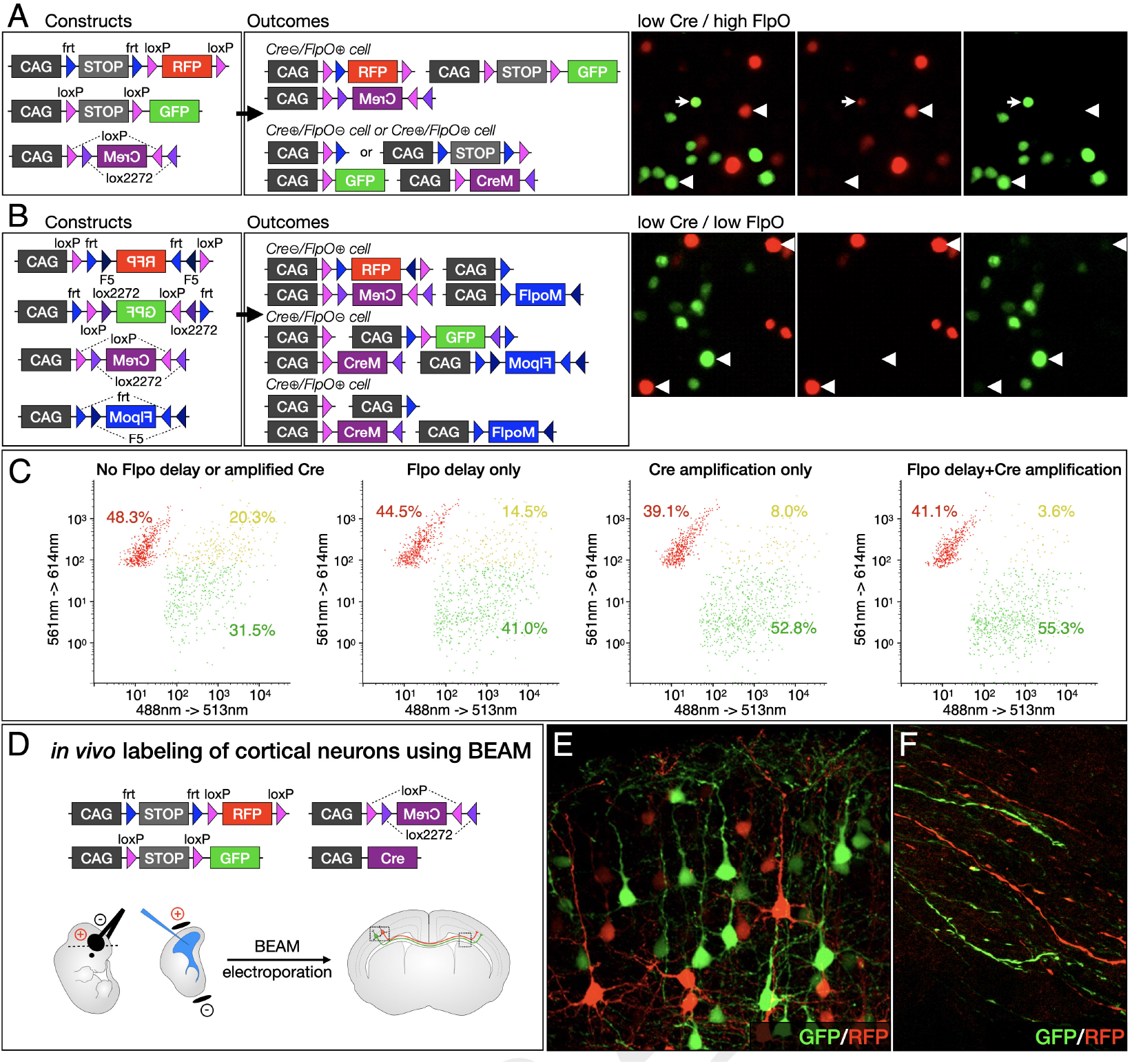
BEAM system results in distinct expression of *GFP* or *RFP*, both *in vitro* by transfection into 293T cells and *in vivo* by *in utero* electroporation into cortical progenitors. (A) Incorporating recombinase-mediated delayed expression of *RFP* and recombinase-driven signal amplification substantially reduces overlapping expression of *GFP* and *RFP* (arrowheads, right). Only rare CRE-positive cells maintain even low-level, residual *RFP* expression (arrows, right). (B) *FLEX* cassettes produce recombination with minimal crossover between fluorophores (arrowheads, right), but recombination is less efficient. (C) Combining delayed *FlpO* expression with *Cre* amplification dramatically reduces overlap between GFP- and RFP-expressing cells (yellow cells in right upper quadrant). (D-F) *In vivo* labeling of cortical neurons using BEAM at E14.5 (D). Electroporated neurons express either GFP or RFP, and migrate to appropriate locations within cortex, residing intermingled with no apparent positional preference (E). Electroporated neurons extend axons across the corpus callosum that are intermingled, but distinctly labeled with either GFP or RFP (F).

We also generated equivalent reagents for mosaic genetic analysis using flip excision configurations, in which the coding sequence for *GFP* or *RFP* is initially oriented in the antisense direction, and is flanked by inverted repeats of incompatible *lox* sites (*FLEX*) or *frt* sites (*FREX*). Therefore, CRE-or FLPO-mediated recombination will reverse the orientation of GFP or RFP, activating expression of the corresponding fluorescent protein (Figure S2A and S2B). Each cassette is additionally flanked by direct repeats of *frt* sites (*frt-FLEX-frt*) or *loxP* sites (*loxP-FREX-loxP*), so that, while one recombinase activates expression, the other recombinase excises the entire cassette, turning off expression. Sparse transfection with *Cre* and *FlpO*, along with the corresponding amplification constructs, results in cells that receive a single recombinase and activate expression of *GFP* or *RFP*, and cells that receive both recombinases and express neither (Figure 2B). These flip excision constructs provide an alternative to transcriptional *STOP* cassettes, which are known to allow low levels of read-through, especially when placed in front of a strong promoter, such as *CAG*. CRE-postive green cells should have no history of any *RFP* expression, and FLPO-positive red cells should have no history of any *GFP* expression. Because the two-step inversion and excision process is substantially less efficient than direct excision(19), little expression of *GFP* or *RFP* develops in double-positive cells. However, for the same reason, lower *GFP* and *RFP* expression levels are present in cells transfected with a single recombinase. In addition, fewer cells are labeled in total, because transfection with each recombinase is sparse, and double positive cells remain unlabeled.

To investigate whether the BEAM plasmid system can also be used successfully *in vivo*, we introduced the constructs into cerebral cortex progenitors in mice by *in utero* electroporation on embryonic day 14.5 (E14.5), and analyzed the resulting brains on postnatal day (P4). Similar to the results obtained *in vitro*, we observed only red cells in the absence of *CAG-Cre*, and increasing numbers of green cells with escalating doses of *CAG-Cre*, enabling the large majority of cells to be labeled green (Figure S3). Importantly, the green (CRE-positive) and red (CRE-negative) neurons both migrate into cortex and reside intermingled with each other, with no indication that one population is otherwise phenotypically different from the other. High-magnification confocal imaging shows that both populations of neurons have normal pyramidal morphologies and long apical dendrites that arborize in layer I (Figure 2A-2B). In addition, both red and green axons extend normally across the corpus callosum (Figure 2C).

### BEAM efficiently reports genomic recombination status

In order to determine how closely GFP and RFP expression track with the recombination status of transfected cells in vivo, we electroporated the BEAM plasmids into the cerebral cortex of *R26*^*loxP(STOP)loxP-LacZ*^ mice (Soriano, 1999) on E14.5. We collected the brains at P7, performed immunocytochemistry for β–GAL, and examined co-expression with GFP and RFP by confocal imaging (Figure 3A-3H). Highly effective BEAM operation would produce few green cells that are β-GAL-negative or red cells that are β-GAL-positive. However, some β-GAL-positive cells that do not express GFP are expected, because the electroporated plasmids are epigenomic, and therefore are gradually diluted in postmitotic neurons after progenitors undergo multiple rounds of mitosis, while the recombined *R26*^*LacZ*^ allele is transmitted together with the rest of the genome during each round of mitosis. Importantly, we find almost perfect co-localization between GFP and β-GAL immunolabeling (91.7±5.5% of GFP-positive cells were also β-GAL-positive), although some β-GAL-positive cells are GFP-negative, as expected. Conversely, there is almost no overlap between RFP and β-GAL immunolabeling (4.0±2.4% of RFP-positive cells were also β-GAL-positive).

**Figure 3.**
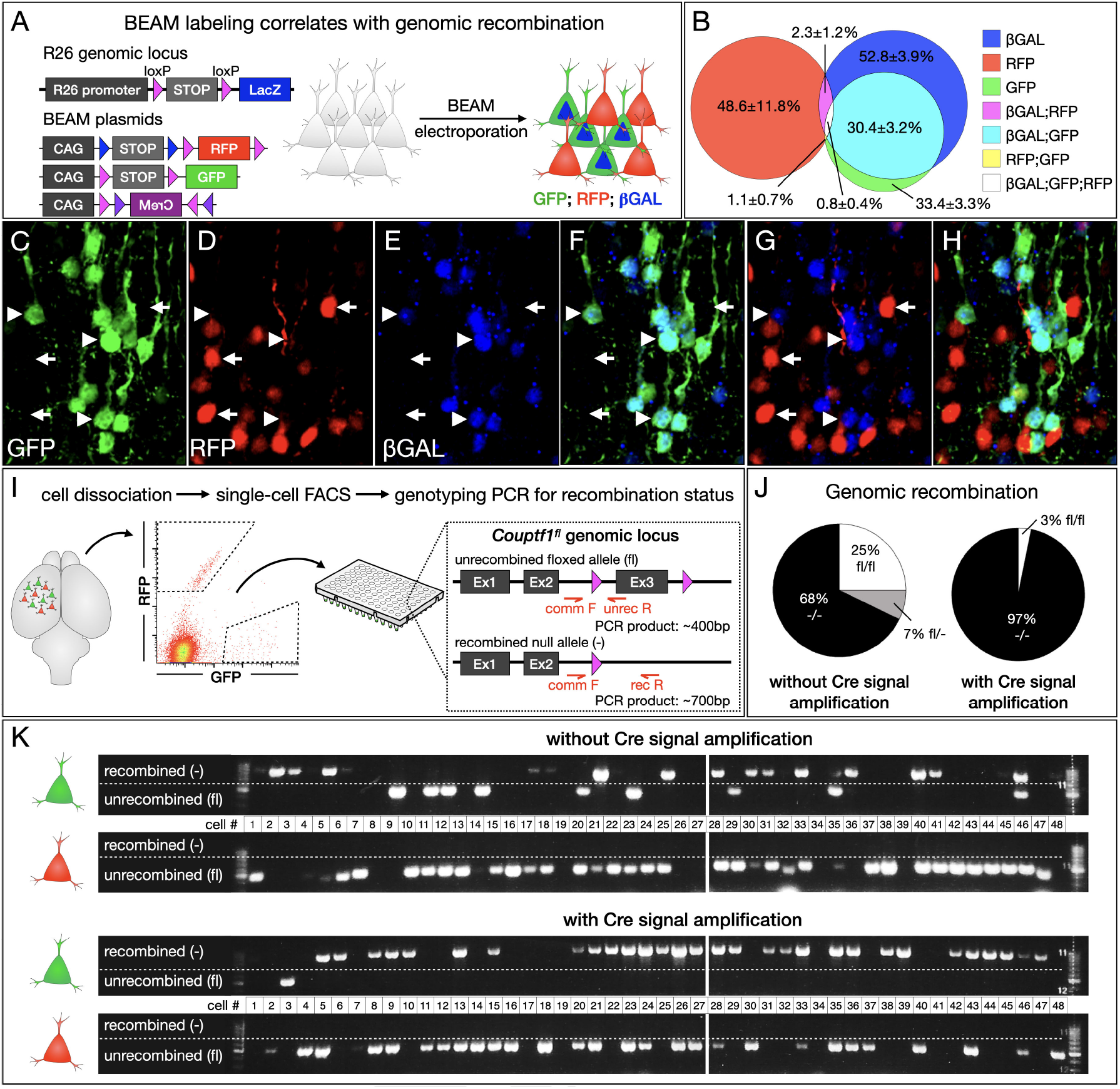
BEAM-driven expression of GFP and RFP accurately reflects genomic recombination status. (A-H) BEAM electroporation into *Rosa26*^*loxP-STOP-loxP-LacZ*^ mice (A). Most GFP-positive neurons are also β-GAL-positive (arrowheads in B-G), while most RFP-positive neurons lack β-GAL staining (arrows in B-G). Quantification (H). (I-K) BEAM labeling efficiently reports gene deletion in *Couptf1*^*fl/fl*^ mice. Electroporated neurons were dissociated, and single cell sorting was performed into a 96-well plate to obtain 48 green cells and 48 red cells (I). Single cell genotyping PCR demonstrates that all tdTomato-positive neurons remain unrecombined. GFP-positive cells were recombined with considerably higher efficiency when *FLEX(CreM)* was used to amplify CRE activity (K). Quantification (J).

We also tested how closely excision of a floxed gene of interest correlates with fluorescent reporter expression, by electroporating BEAM plasmids into the cerebral cortex of *Couptf1*^*fl/fl*^ mice(20)(21) and analyzing the recombination status of GFP and RFP labeled cells (Figure 3I). Electroporations were performed at E14.5, tissue was collected at P4, dissociated, and individual neurons were then FACS purified into 96-well plates. Single-cell PCR was used to interrogate the recombination status of the *Couptf1* locus in each isolated cell. Strikingly, we find that 96% of BEAM-electroporated green cells (31/32 cells) were fully recombined when recombinase amplification was used, while only 68% (19/28 cells) were recombined when the *CAG-FLEX(CreM)* plasmid was omitted, with 7/9 cells genotyping as fully unrecombined and 2/9 genotyping as heterozygous. Importantly, all of the genotyped BEAM-electroporated red cells remained unrecombined (64/64 cells), indicating that there is no spontaneous CRE activity resulting from the presence of *CAG-FLEX-CreM*. Taken together, these data indicate that BEAM plasmids can be effectively used to investigate gene function by electroporation into genetically modified floxed mice.

### Cell autonomous gene function can be rigorously investigated using BEAM

As discussed above, dual-population mosaic analysis is a particularly powerful approach for investigating the cell autonomy of a phenotype. In the BEAM system, RFP-labeled (CRE-negative) cells provide an internal and spatially interspersed control for direct comparison to experimental GFP-labeled (CRE-positive) cells within the same animal, eliminating pleiotropic effects resulting from potential abnormalities in other mutant cell populations. As a proof-of-principle experiment, we re-investigated the effect of *Satb2* loss-of-function on the trajectory of callosal projection neuron (CPN) axons in *Satb2*^*fl/fl*^ mice(22)(23)(24). In *Satb2*^*-/-*^ mice, superficial-layer neuron axons fail to project toward the corpus callosum, and project subcortically instead(25)(26). In addition, these neurons abnormally activate expression of *Ctip2*, a transcriptional control normally specific to corticofugal projection neurons (though present later in interneurons at lower expression levels), and that is critical for their development(27)(25). Previous studies have proposes that these defects in axonal targeting are due to a cell-autonomous failure of callosal projection neuron subtype differentiation, and exclude a number of alternative possibilities, including abnormalities in glial structures necessary for midline fusion(25).

BEAM plasmids were electroporated into the cerebral cortex of *Satb2*^*fl/fl*^ mice at E14.5, and the resulting brains were examined at P7. We find that a number of GFP-labeled neurons undergo migrational arrest in the white matter (Figure S4B and S4F), their axons enter the internal capsule and fasciculate with other corticofugal axons (Figure S4C and S4G), and project into the thalamus and cerebral peduncle (Figure 4C-D and 4G-H). Immediately adjacent RFP-positive control neurons migrate normally, and do not project axons into the internal capsule. These results are similar to what has been observed in *Fezf2* and *Ctip2* overexpression experiments(28)(29), and reinforce that abnormal migration in *Satb2*^*-/-*^ mice might result, at least in part, from aberrant and premature expression of *Ctip2* and other molecular controls of subcerebral projection neuron differentiation, as previously proposed(28)(25). Importantly, in brains electroporated with BEAM plasmids carrying nuclear targeted RFP and GFP, immunostaining for SATB2 and CTIP2 demonstrates that almost all tdTomato-labeled neurons are SATB2-positive and CTIP2-negative (Figure 4G). Conversely, a large majority of GFP-labeled neurons lack SATB2 and have up-regulated CTIP2 (Figure 4H). One surprising finding from these experiments is the large number of GFP-positive axons that project successfully across the corpus callosum in *Satb2*^*fl/fl*^ brains (Figure S4D and S4H). These data sharply contrast with the absolute failure of cortical neurons to project across the corpus callosum in *Satb2*^*-/-*^ mice(25). Although recombination at the Satb2 locus is expected to fail in a small percentage of GFP-labeled neurons, these few escapers cannot account for the largely unchanged proportion of GFP-labeled axons projecting across the corpus callosum. Therefore, it seems likely that axons of GFP-labeled *Satb2*^*-/-*^ neurons, though impaired, are often able to fasciculate with axons of unrecombined *Satb2*^*fl/fl*^ neurons (both RFP-positive and unelectroporated) to project successfully across the corpus callosum. These results further highlight the advantages of such genetic mosaic analysis via BEAM in elucidating both cell-autonomous and non-cell-autonomous, as well as other nuanced and relative or absolute disruptions of cellular function.

**Figure 4.**
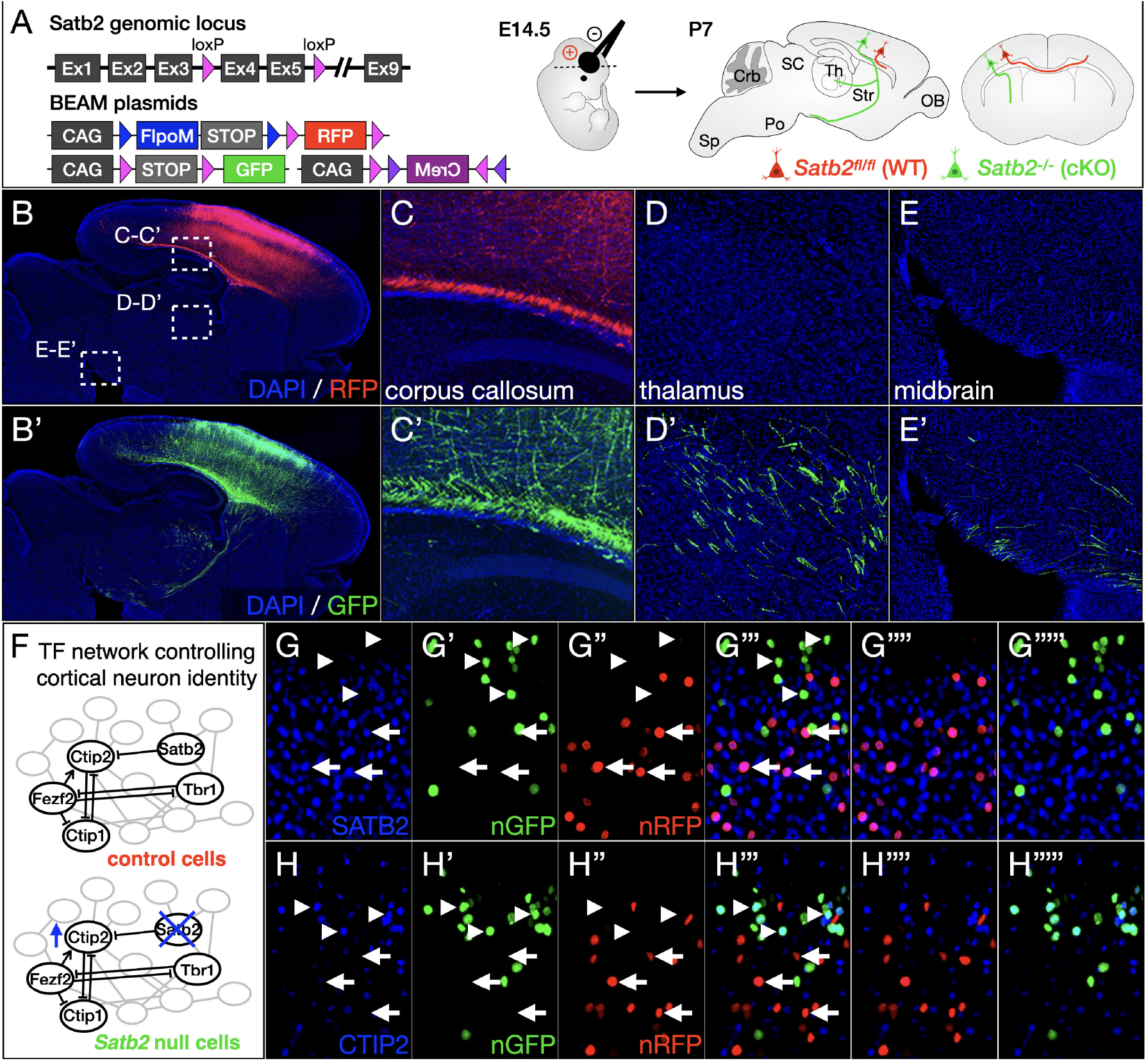
BEAM enables mosaic gene deletion and investigation of cell-autonomous gene function in conditional null mice. (A-E) Electroporation of BEAM into *Satb2*^*fl/fl*^ mice demonstrates abnormal axon targeting by conditional null neurons (A). Although both GFP and RFP-positive axons are present in the corpus callosum (C-C’), only GFP-positive axons are redirected to the thalamus (D-D’) and midbrain (E-E’). (F-H) Electroporation of nuclear BEAM into *Satb2*^*fl/fl*^ mice results in aberrant expression of CTIP2 by conditional null neurons (F). Immunolabeling for SATB2 and confocal imaging demonstrates that GFP-positive neurons do not express SATB2 (arrowheads, G-G”‘), while RFP-positive neurons do (arrows). Conversely, immunolabeling for CTIP2 is absent from RFP-positive neurons (arrows, H-H”‘), but is present in GFP-positive cells (arrowheads).

### Rapid screening of gene function using BEAM

Internally controlled experiments provide a particularly attractive platform for functional screening of genes predicted to regulate a biological process of interest. In order for such an approach to be optimally versatile, it should not rely solely on the use of genetically-modified conditional knockout mice. We therefore sought to determine whether BEAM would be compatible with existing gain-and loss-of-function strategies that rely exclusively on delivery of exogenous DNA into wild-type mice by transfection or transduction.

The most straightforward and broadly used gain-of-function strategy is to mis-express a gene of interest in a population of cells from which it is normally absent, using a strong ubiquitous promoter. As proof-of-principle, we generated a *CAG-*driven *FLEX* construct for the transcription factor *Fezf2*, which is normally expressed during cortical development by early-born corticofugal projection neurons that extend axons to the thalamus and brainstem, but excluded from late-born callosal projection neurons that extend axons across the corpus callosum to the contralateral hemisphere(28)(30)(31). Misexpression of *Fezf2* by late-born callosal projection neurons causes them to redirect their axons to the thalamus and brainstem(28). Co-electroporating the *CAG-FLEX-Fezf2* construct, along with the previously described BEAM plasmids, should result in control RFP-labeled cells in which *Fezf2* remains in the antisense orientation, and GFP-labeled cells in which CRE activates expression of *Fezf2* (Figure 5A). Strikingly, while control RFP-labeled axons project exclusively across the corpus callosum, many *Fezf2*-overexpressing GFP-labeled axons are misrouted to the thalamus and brainstem (Figure 5B-5E), rigorously confirming the prior results.

**Figure 5.**
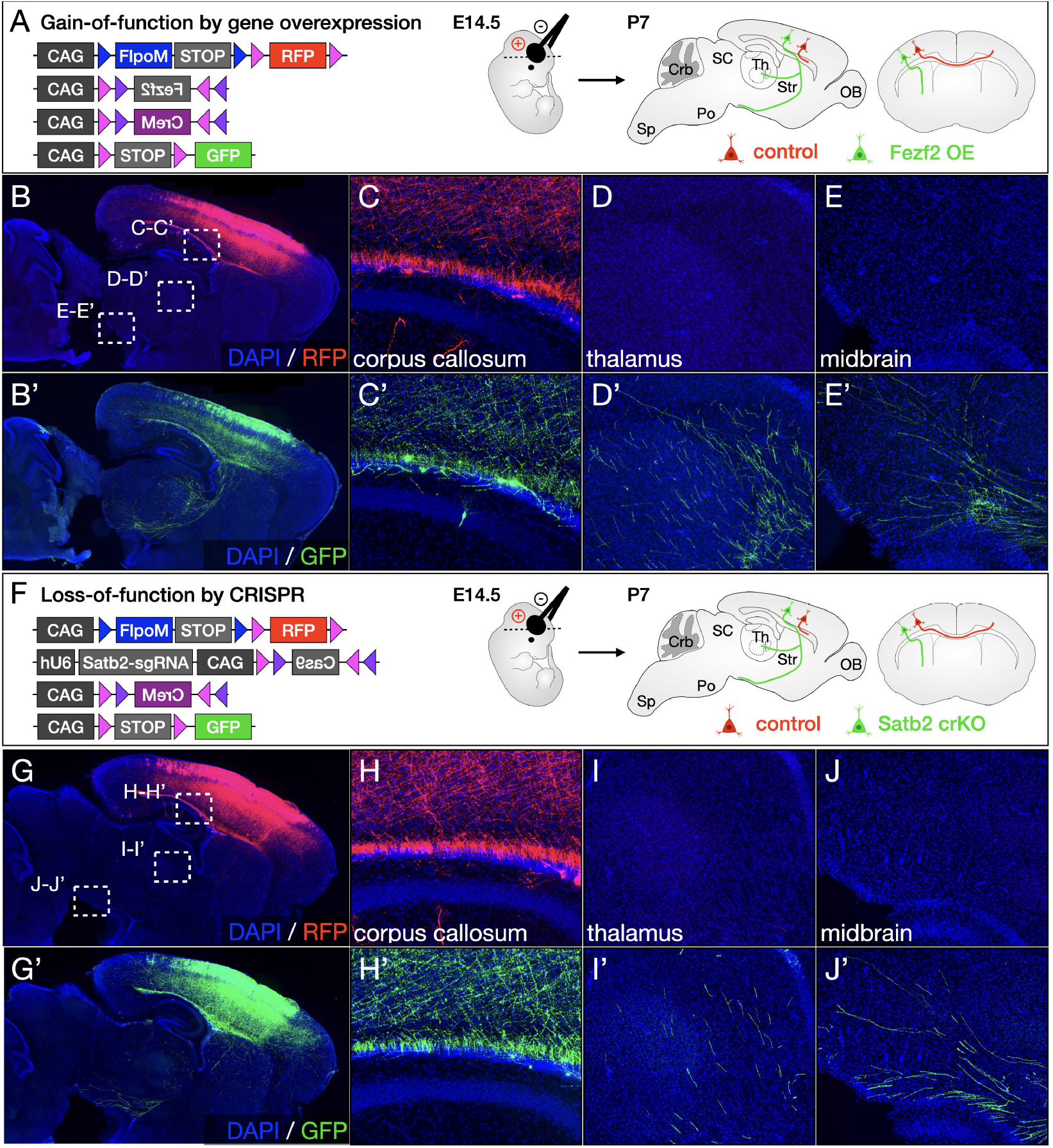
BEAM provides a platform for rapid in vivo screening of gene function by gain- and loss-of-function approaches. (A-E) Mis-expression of *Fezf2* in callosal projection neurons using BEAM (A). CRE-positive GFP-labeled neurons ectopically express *Fezf2* due to activation of the *CAG-FLEX(Fezf2)* construct, causing them to extend aberrant projections to the thalamus and brainstem (D’ and E’) instead of the contralateral hemisphere (C’). Intermingled control RFP-labeled cells project exclusively across the corpus callosum (D-E). (F-J) CRISPR-Cas9 ablation of *Satb2* in callosal projection neurons using BEAM (F). Although all neurons express guideRNAs targeting the *Satb2* gene, *Cas9* expression from a *CAG-FLEX(Cas9)* construct is activated by CRE exclusively in GFP-labeled cells, introducing mutations that inactivate *Satb2* function and redirecting their axons from the corpus callosum (H’) to the thalamus and brainstem (H’ and I’). In the absence of *Cas9* expression, intermingled control RFP-labeled neurons are unaffected by the guideRNA, and project normally across the corpus callosum (H-J).

For proof-of-principle genetic loss-of-function experiments, we focused on CRISPR-Cas9-mediated gene editing. Among the wide range of tools currently available for loss-of-function, we predicted that this approach would be more sensitive for cell-autonomous processes because homozygous indels generally result in complete loss of gene function. We generated a bicistronic construct with a U6 promoter driving expression of a guide RNA, and a CAG promoter driving expression of Cas9 in a CRE-dependent manner using *FLEX* (Figure 5F). To enable direct comparison to the loss-of-function experiments with gene deletion using a *floxed* allele, we designed guide RNAs targeting the coding sequence of *Satb2*, and electroporated these constructs into developing cortex of wild-type mice at E14.5. Efficiency of CAS9-mediated gene deletion was examined by SATB2 immunocytochemistry. We find that GFP-labeled neurons expressing *Cas9* are predominantly SATB2 negative by immunocytochemistry, while control RFP-labeled neurons are predominantly SATB2 positive (Figure S5C). Confirming previously reported results(32), CAS9-mediated ablation of *Satb2* expression is correlated with abnormal axon trajectories, with many GFP-labeled axons aberrantly projecting to the thalamus and brainstem (Figure 5I-5J), while RFP-labeled axons project exclusively across the corpus callosum (Figure 5H). These results again rigorously confirm the prior reports, with BEAM’s genetic mosaic analysis highlighting the cell-autonomy of *Satb2* function.

### BEAM analysis can rigorously and unequivocally identify abnormalities in the timing of developmental processes

In addition to providing a rigorous experimental approach for investigating cell autonomy, dual-population mosaic analysis is particularly well-suited for uncovering phenotypes that are subject to substantial inter-trial variability, such as the timing of developmental events. To illustrate this application, we used BEAM to investigate regulation of neuronal migration by *Satb2* in the developing cerebral cortex. Superficial-layer neuron migration is severely disrupted in *Sabt2*^*-/-*^ mice, and few neurons generated at later stages of cortical development are able to enter the cortical plate and reach their appropriate laminar position(25)(26). We electroporated BEAM plasmids carrying nuclear targeted RFP and GFP into cortical progenitor cells at E14.5, and collected brains for analysis 3 days later, at E17.5. In wild-type embryos, both GFP- and RFP-labeled neurons migrate efficiently into the cortical plate (Figure 6B). In *Satb2*^*fl/fl*^ embryos, control RFP-labeled neurons still reach the cortical plate normally, but *Satb2* null GFP-labeled neurons become stalled in the progenitor and intermediate zones, with few reaching the cortical plate (Figure 6C). To investigate whether this represents a delay or a complete failure of migration, we repeated the same experiments, but analyzed the brains at P7. Interestingly, although a subset of Satb2 null GFP-labeled neurons undergo migrational arrest in the white matter and never enter the cortical plate, the majority of them join control tdTomato-labeled neurons in layer II/III of cortex, and in fact migrate past them to reside in aberrantly superficial positions (Figure S6A-S6B). These direct wild-type and mutant population comparisons of interspersed neurons within the same areas of the same brains enable identification of even subtle dynamic alterations during development and maturation.

**Figure 6.**
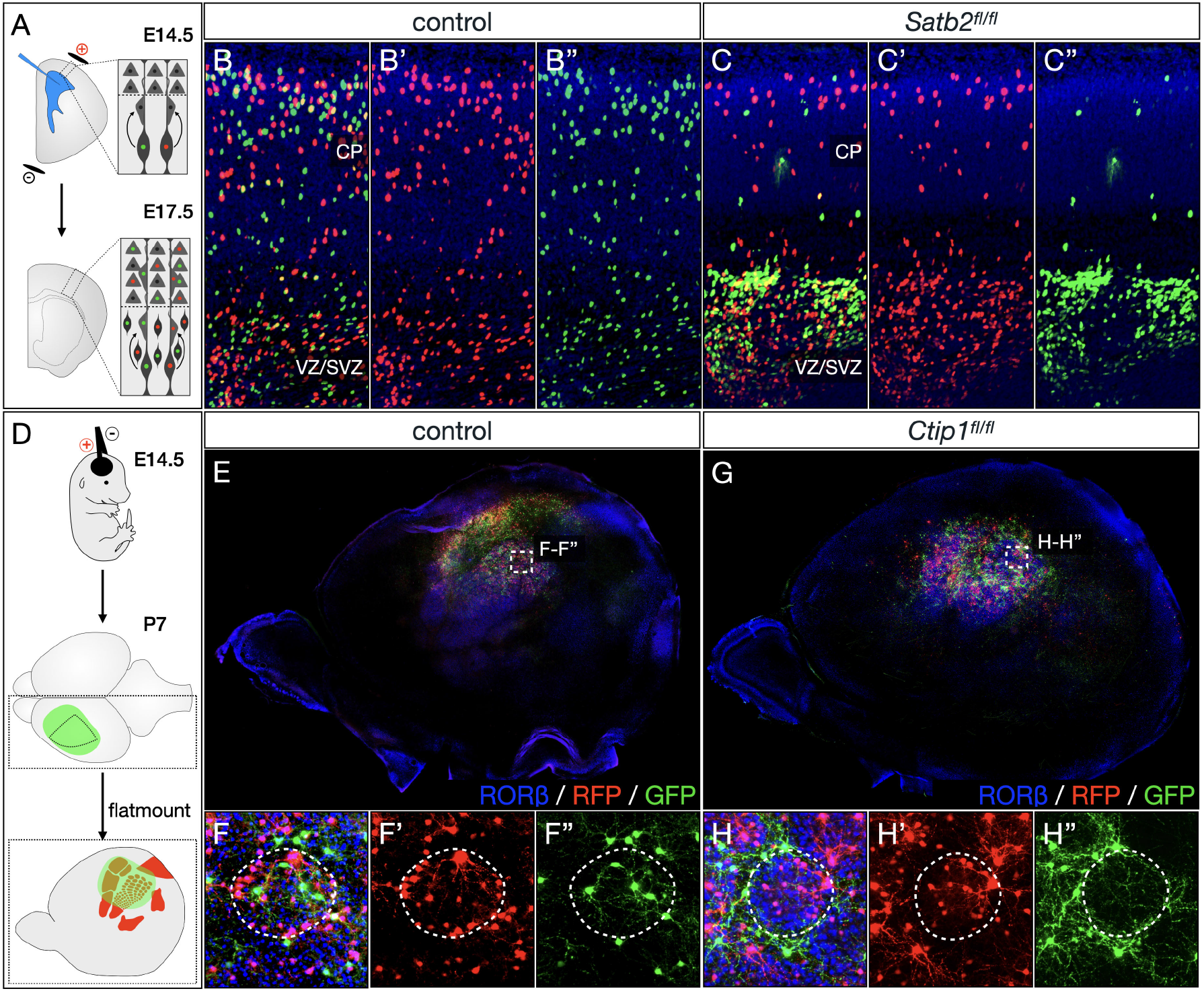
BEAM readily identifies abnormal timing of developmental processes, shifts in the spatial distribution of cells. (A-C) BEAM reveals cell autonomous migrational delay of cortical neurons in the absence of *Sabt2* function. While both RFP-positive and GFP-positive neurons migrate into cortex at E17.5 following E14.5 electroporation in wild-type mice (B-B”), only RFP-positive control neurons migrate successfully into cortex in *Satb2*^*fl/fl*^ mice (C-C”). GFP-positive mutant neurons are cell-autonomously impaired in their migration in *Satb2*^*fl/fl*^ mice (B”). (D-H) BEAM demonstrates abnormal spatial distribution of neurons lacking *Ctip1* within the barrel field. Following electroporation at E14.5 both RFP- and GFP-labeled neurons integrate into barrels by P7 in wild-type mice, extending dendrites within the barrel to receive input from thalamocortical axons (E-F). In *Ctip1*^*fl/fl*^ mice, control RFP-labeled neurons adopt this normal configuration, while *Ctip1* null GFP-positive neurons and their dendrites are strikingly excluded from barrels (G-H).

We further applied BEAM to investigate the extension of callosal projection neuron axons across the midline and into the contralateral hemisphere, which is delayed in the absence of *Couptf1* function(33). BEAM plasmids were electroporated at E14.5, and brains were collected for analysis at P0, when the axons of late-born superficial layer callosal projection neurons are still en route to the contralateral hemisphere. GFP-labeled axons behaved identically to control RFP-labeled axons in wild-type brains, crossing the corpus callosum and invading the contralateral hemisphere. In *Couptf1*^*fl/fl*^ brains, however, GFP-labeled mutant axons reached corpus callosum, but were absent from the contralateral hemisphere (Figure S6C-S6H). The presence of an internal control of fully interspersed neurons in the same brains crisply rules out the possibility of even subtle mismatched developmental stages across genotypes.

Another broad category of phenotypes that can be investigated extremely effectively using BEAM are those relating to the spatial distribution of cells or cellular processes. As proof-of-principle, we used BEAM to examine the organization of layer IV neurons in somatosensory cortex of *Ctip1*^*fl/fl*^ mice. In the vibrissal barrel field, thalamocortical input from each whisker condenses into distinct clusters, each relaying information from a single whisker to a cytoarchitecturally distinct cluster of layer IV neurons known as a barrel. *Ctip1* is necessary for proper differentiation of layer IV neurons, and their ability to organize into barrels is severely impaired in its absence(34). Electroporation of BEAM into wild-type cortex at E14.5 results in both RFP- and GFP-labeled neurons adopting an unbiased distribution intermingled across barrel cortex, with higher density of neurons in the barrel walls, and most dendrites extending toward the center of each barrel (Figure 6E-6F). Electroporation of BEAM into *Ctip1*^*fl/fl*^ brains leads to a similar overall distribution of control RFP-labeled cells. Strikingly, however, *Ctip1* conditional null GFP-labeled cells and their dendrites position themselves preferentially between barrels, segregating themselves in the septae (Figure 6F-6H). The mosaic results are sharply distinguished within the organization of even individual barrels.

### BEAM controls for procedural variability enabling efficient investigation of cell proliferation and survival

Experiments to investigate cellular proliferation and survival can be difficult to interpret due to significant variability in the number of cells labeled across trials. To investigate whether BEAM can provide more robust measures of these phenotypes by enabling normalization across trials, we performed experiments involving manipulation of cortical progenitor proliferation. Previous studies have shown that the *Wnt* signaling pathway regulates the cell cycle in cortical progenitors, so that overexpression of β*-catenin* stimulates proliferation of cortical progenitors(35), while a dominant negative *Tcf4* (*Tcf4-DN*) inhibits proliferation(36). We generated *CAG*-driven *FLEX* constructs for expression of β*-catenin-2A-H2B-EGFP* or *Tcf4-DN-2A-H2B-EGFP* (Figure 7A). Using BEAM, we find a 25% ± 9% increase in the ratio of GFP:RFP labeled cells in β-catenin electroporations relative to control experiments (Figure 7C-7D, quantification 7B), indicating increased proliferation or survival of cells overexpressing β*-catenin*. Conversely, the ratio of GFP:RFP labeled cells in *Tcf4-DN* electroporations is decreased by 62% ± 15% relative to control experiments (Figure 7D-7E, quantification 7B), indicating decreased proliferation or survival of cells overexpressing *Tcf4-DN*. Importantly, without normalization, the trends toward more cells with overexpression of β*-catenin* and fewer cells with overexpression of *Tcf4-DN* are still present, but the differences across conditions are no longer statistically significant. Normalization to an internal control eliminates an important source of experimental variability, substantially increasing statistical power enabling smaller changes to be detected without a large sample size.

**Figure 7.**
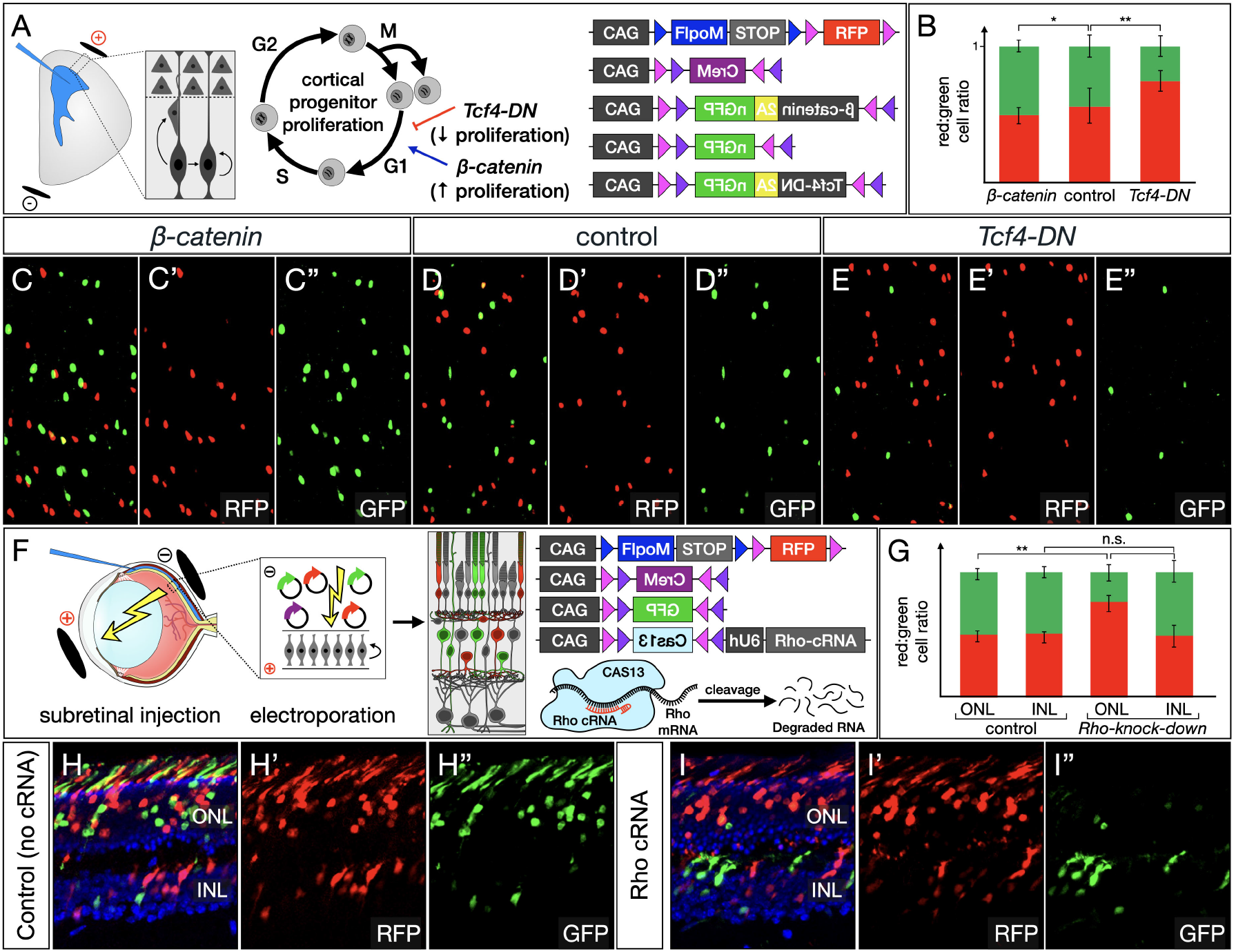
BEAM enables unequivocal investigation of changes in mitotic activity and cell survival. (A-E) BEAM facilitates studies of cell cycle dynamics in cortical progenitors (A). The relative ratios of control RFP- and genetically manipulated GFP-labeled cells after electroporation at E12.5 and analysis at E17.5 reflect the relative balance of proliferation and cell death in each subset. Compared with control experiments (D-D”), *b-catenin* overexpression increases the relative number of GFP-labeled cells (C-C”), while a dominant negative *Tcf4* mutant decreases the relative number of GFP-labeled cells (E-E”). Quantification (B). (F-I) BEAM reveals selective cell death due to photoreceptor degeneration (F). Approximately equal numbers of RFP- and GFP-positive cells are present in control retinas both in the INL and ONL. CAS13-mediated knockdown of *Rho*, does not affect survival of cells in the INL, however there is a striking decrease in GFP-positive cells in the ONL, compatible with photoreceptor degeneration and death in the absence of *Rho*. Note also the complete absence of GFP-positive outer segments in *Rho* knockdown (I”), while there is robust GFP-labeling of outer segments in cotrol experiments (H”). Quantification (G).

Procedural variability in the number of cells labeled across experiments also complicates efforts to investigate cell survival. Here, we demonstrate that CAS13-mediated knockdown can be used to recapitulate a neurodegeneration phenotype in the retina (Figure 7F). We focused on retinitis pigmentosa, the most common mendelian degenerative retinopathy, which is caused by mutations in one or more genes important for photoreceptor development and function leading to progressive loss of rods and decreased vision. Rhodopsin null mice were one of the earliest published mouse models of retinitis pigmentosa(37). In *Rho*^*-/-*^ mice, rods fail to elaborate outer segments during development and most photoreceptors are lost by 3 months of age. We first tested guide RNAs to determine if we could induce efficient kockdown of *Rho* and find that while there is extensive overlap of RHO immunostaining with the outer segments of electroporated photoreceptors in control experiments, there is a complete lack of overlap when guide RNAS targeting *Rho* are used, indicating efficient knockdown (Figure S7). In BEAM experiments, we find approximately equal ratios of RFP- and GFP-positive cells in both the inner nuclear layer (INL) where electroporated bipolar, Müller glia and amacrine cells reside, and in the outer nuclear layer (ONL) where photoreceptors reside (1 and 0.94, no statistically significant difference; Figure 7H, quantification 7G). Although in Rho knockdown experiments the ratio of RFP-to GFP-positive cells remains approximately equal in the INL (1.09, no statistically significant difference compared to control INL; Figure 7I, quantification 7G), the ratio is strikingly changed in favor of RFP-positive cells (3.37, p<0.01; Figure 7I, quantification 7G). Importantly, the few remaining GFP-positive cells in the ONL generally exhibit low levels of fluorescence and no GFP-positive outer segments can be discerned in *Rho* knockdown retinas. These results are consistent with death of photoreceptors due to loss of *Rho* expression, and together with our experiments manipulating the cell cycle dynamics of cortical progenitors illustrate the power of mosaic analysis using BEAM to investigate cell proliferation and survival.

### Evaluating the protomap hypothesis of cortical area development at the level of individual radial units using BEAM

The protomap hypothesis of cortical development proposes that area identity is specified in cortical progenitors, and that information is transferred to postmitotic neurons in each radial unit(38)(39). Manipulating morphogen activity or transcription factor function is sufficient to change the relative size and position of cortical areas(40)(21)(41). For instance, loss of *Couptf1* function throughout the cerebral cortex results in a dramatic expansion of motor areas, and a reciprocal reduction in the size of sensory areas (Figure 8A). However, it is not known whether area identity is independently programmed within each radial unit, or is interdependent across adjacent radial units. To investigate these possibilities, we performed BEAM electroporations into *Couptf1*^*fl/fl*^ brains, so that a subset of CRE-negative progenitors gave rise to RFP-labeled control neurons and a subset of CRE-positive progenitors gave rise to GFP-labeled *Couptf1* null neurons. We then examined area-specific output connectivity and gene expression of electroporated neurons.

**Figure 8.**
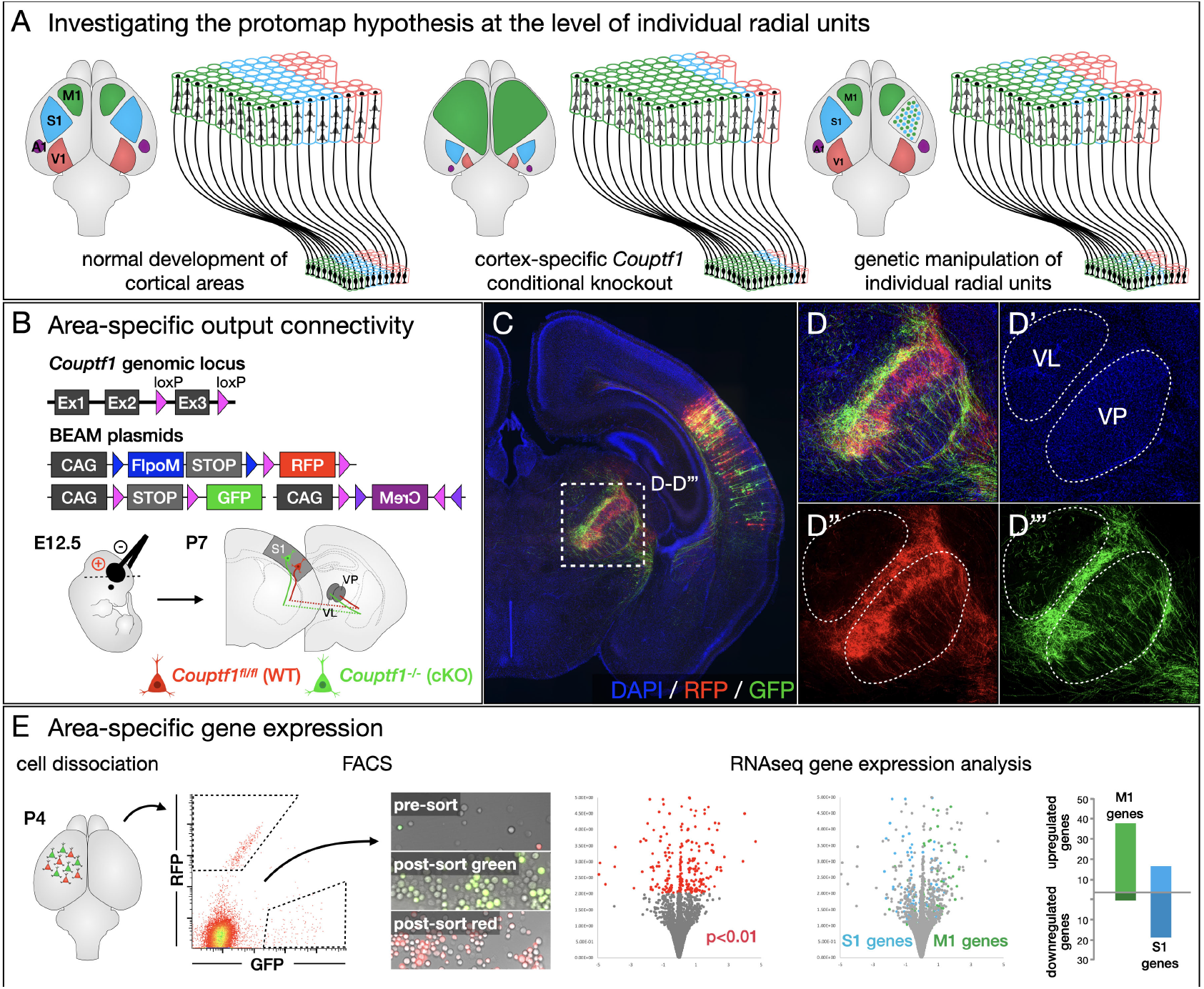
Investigating the protomap hypothesis at the level of individual radial units reveals that area identity is independently specified in each progenitor cell and its progeny. (A) During normal development, individual cortical progenitors acquire specific area identities, and transfer this information to postmitotic neurons generated by each radial unit. Cortex-wide loss of transcription factors that specify area identity can lead to relative changes in the size and position of cortical areas. It is not known whether area specification occurs independently within each radial unit, or whether there are mechanisms driving interdependent area specification of adjacent radial units. (B-D) BEAM electroporation into *Couptf1*^*fl/fl*^ mice reveals that area-specific output connectivity is independently specified in adjacent radial units. Each cortical area establishes axonal projections to specific thalamic nuclei, with terminal axonal fields in each nucleus labeled by electroporation at E12.5 (B). While control RFP-labeled axons originating from somatosensory radial units project primarily to the ventral posterior (VP) sensory nucleus (D”), GFP-labeled axons originating from intermingled *Couptf1* null radial units avoid the VP nucleus, with aberrant increased density in the immediately adjacent ventral lateral (VL) motor nucleus (D”‘). (E) RNAseq analysis of BEAM-electroporated *Couptf1*^*fl/fl*^ neurons demonstrates that expression of genes normally expressed in motor vs. sensory cortex is regulated independently in adjacent radial units. *Couptf1*^*fl/fl*^ brains were electroporated at E12.5, then cortical tissue was collected from somatosensory cortex at P4 and dissociated. Control RFP-labeled cells and conditional null GFP-labeled cells were purified by FACS, and gene expression was analyzed by RNAseq. Among differentially expressed genes, motor-specific genes were upregulated, while sensory-specific genes were downregulated.

Each cortical area establishes output connectivity to a specific thalamic nucleus, with somatosensory cortex projecting mainly to the ventral posterior nucleus (VP), and motor cortex projecting primarily to the ventral anterior and ventral lateral nuclei (VA and VL). We electroporated BEAM into somatosensory cortex of wild-type and *Couptf1*^*fl/fl*^ at E12.5, when corticothalamic projection neurons are being generated. In control electroporations, both RFP-and GFP-labeled axons project to VP, and arborize with a similar distribution (Figure S8A-8C). Electroporation into *Couptf1*^*fl/fl*^ brains also resulted in RFP-labeled axons that project and arborize primarily within VP, while GFP-labeled axons arising from intermingled *Couptf1* conditional null radial units largely avoid arborizing within VP, and shift toward arborizing in VL instead (Figure 8B-8D).

Another feature that defines each cortical area is gene expression. In parallel experiments, we microdissected BEAM electroporated *Couptf1*^*fl/fl*^ somatosensory cortex at P3, dissociated the tissue, and FACS purified RFP- and GFP-labeled cells arising from intermingled radial units in the same brain. We then performed RNAseq gene expression analysis, and identified differentially expressed transcripts (Figure 8E). By using previously published datasets of motor- and somatosensory-specific genes(34), we identified that 74 of 201 differentially expressed genes in these BEAM *Couptf1* loss-of-function experiments are area-specific, with overall upregulation of motor-specific genes and downregulation of sensory-specific genes. Taken together, these data indicate that area-specific output connectivity and gene expression are independently determined in neighboring radial units, simultaneously grounding results in prior work via the tdTomato wt neurons, while discovering the cell-autonomous, individual radial unit basis of areal development via fully interspersed GFP mutant neurons in the same brains.

These results highlight that BEAM not only flexibly enables new investigations with added discovery power, rigor, ease, and comparative insight, but also enables elucidation of prior work and observations at more detailed levels of mechanism and cell autonomy. Even subtle effects from molecular and cellular manipulations are placed in stark contrast by fully intermingled cells in exactly the same areas of the same brains– or, by extension, of any cellular, tissue, or organ system under investigation.

## Discussion

The BEAM genetic mosaic system presented here offers a flexible and powerfully enabling approach for molecular functional analyses that relies on combinatorial recombinase activation to generate two genetically distinct, fully interspersed, fluorescently-labeled populations of cells for comparative analysis in the same spatial domain of the same tissue and animals simultaneously. BEAM offers critical advantages over genome-based tools, such as MADM and MASTR, as well as over existing plasmid-based tools, such as Double-up and Beatrix. BEAM enables sharper and earlier delineation of control and experimental cells, it allows for a broad range of experimental manipulations (including floxed alleles, overexpression, Cas9-mediated knockout), and it has been validated across multiple DNA delivery strategies (chemical and physical transfection, as well as viral transduction).

### Strategies for binary gene expression by exogenous DNA delivery

Accomplishing binary gene expression and fluorescent labeling of cells by asymmetric recombinase activity in the context of plasmid transfection or viral transduction presents a number of challenges. Delivery of a single recombinase, such as CRE, to a subset of transfected cells is sufficient to activate expression of one set of genes and inactivate expression of another. However, there is a temporal window before CRE becomes functional, during which control cells can express genes that are destined for later inactivation, and are therefore inconsistent with the final binary outcome. This can lead to control cells with intermediate phenotypes, thus obscuring the full effect of the experimental manipulation.

To overcome this limitation, we designed a second, orthogonal recombinase system to delay expression of these genes. An frt-flanked transcriptional STOP cassette is introduced genetically “in front of” genes intended for CRE inactivation, so that recombination by FLPO is required to activate their expression. With these design elements, there is simultaneous competition between activation by FLPO and inactivation by CRE, very substantially reducing transient expression of these genes in CRE-positive cells. We also generated a version in which an intron interrupted *FlpoM* is incorporated into the delayed expression plasmid, which self-excises to allow expression of a downstream gene. These constructs might be useful in other contexts in which delayed expression is desirable. Moreover, cloning a gene in tandem with the self-excising recombinase for polycistronic expression would enable a short burst of expression of a gene of interest until it is excised along with the recombinase. These flexible approaches offer multiple options to prevent toxicity and other deleterious effects that sometimes arise from expression of genes at high levels or for prolonged periods.

A second challenge to accomplishing binary gene expression arises from incomplete recombination of the multiple copies of reporter DNA in each cell. Because a large number of copies (>100 on average) are delivered to each cell by transfection or AAV-mediated transduction, even a highly efficient recombinase will leave multiple copies unrecombined. This problem is compounded in actively dividing cells, because DNA is randomly distributed with each round of mitosis, so that not all daughter cells will inherit copies of *Cre* recombinase DNA when relatively few copies are initially present.

Here, we have mitigated incomplete recombination by using a FLEX system in combination with an intron-interrupted *Cre* to enable amplification of CRE activity. Importantly, we demonstrate that CRE amplification substantially improves the concordance of fluorescent reporter expression and genomic recombination status, investigated by genotyping of individual FACS purified cells. Moreover, this system exhibits no auto-activation in the absence of a second source of CRE, ensuring that control cells remain CRE-negative throughout the experiment. Although a small number of βGAL-positive;RFP-labeled cells were observed following electroporation into *R26*^*loxP(STOP)loxP-LacZ*^ mice, the complete absence of Tomato-labeled cells with abnormal phenotypes across our experiments conclusively validates the use of the BEAM system for analysis of gene function. As just one striking example of the specificity of the BEAM system, no GFP-labeled control axons project subcortically following mosaic *Fezf2* overexpression or *Satb2* loss-of-function by excision of a floxed allele or CAS9-mediated knock out. It is thus likely that the rare red βGAL-positive;RFP-labeled cells we observed represent non-specific staining or bleed-through of very bright RFP fluorescence into the far-red imaging channel, despite stringent bandpass settings for spectral confocal imaging. This level of essentially absolute contrast between control and manipulated cell populations importantly adds analytic and statistical power to experiments employing BEAM, enabling interrogation of even exquisitely subtle effects, or those that regulate highly refined aspects of neuronal, other cellular, or intercelluar development, organization, or function.

### BEAM enables improved reproducibility of genetic manipulation experiments

Reproducibility and replicability are critical for optimal, rigorous interpretation of experimental data and overall studies, and have been an area of significant interest and concern for the scientific community for many years. More recently, the U.S. National Institutes of Health (NIH) has developed action plans to address these concerns. As one notable example, studies of spinal cord injury have suffered from a lack of replicability, a problem brought into sharp focus by the NINDS-sponsored FORE-SCI project(42). One likely explanation for failures to replicate across studies, as well as the resultant often erroneous conclusions, is that the severity of experimentally-induced injuries and/or level of genetic or viral manipulation varies significantly across trials, and has been found to often lack reproducibility within individual experiments. The ability to intersperse both experimental and control samples within each individually manipulated animal enables control for variable injury, genetic or viral manipulation, timing, and other experimental variables across multiple animals, regions, and other experimental replicates. Although we focus here on developmental phenotypes, we demonstrate that application of BEAM for internally controlled experiments substantially reduces procedural variability, significantly enhancing ability to detect or rule out genuine effects. We also perform proof-of-principle experiments demonstrating that the BEAM system is compatible with AAV-mediated gene transduction, widely used approaches for cell labeling and gene delivery. Therefore, application of BEAM for individually and internally controlled experiments offers substantial potential to enable more rigorous evaluation of potential effects of manipulations, e.g. therapeutic benefits in spinal cord injury regeneration studies, and to increase reproducibility more broadly across a variety of fields.

### Limitations of gene function analysis using BEAM

Although mosaic analysis eliminates multiple sources of experimental variability, it is important to remain mindful of biases inherent to generation of genetically distinct populations of cells using the BEAM recombinase system. Whether an individual cell becomes a control cell or an experimental cell depends on the presence or absence of *Cre*, which is a stochastic event on a cell-by-cell level. However, on a population level, cells that receive a higher copy number of plasmids or viral genomes are also more likely to receive a copy of *Cre*, which is introduced at a lower dose. The average number of plasmids or viral genomes present in an experimental CRE-positive cell is therefore higher than the average number present in a control CRE-negative cell (Figure 1A). Which cells take up a higher copy number of plasmids or viral genomes theoretically might not be entirely random. For instance, some cortical progenitor subtypes might be electroporated more efficiently, based on proximity to the lateral ventricle, cell morphology or expression of specific cell membrane proteins. Similarly, viral transduction efficiency varies widely across different neuron and glial cell types, as determined by the tropism of particular viral serotypes. In addition, cells carrying higher copy numbers of plasmid or viral genomes might respond to the additional foreign DNA by more strongly activating immune cell signaling pathways. With these considerations in mind, we recommend that before transitioning to internally controlled experiments, investigators should first establish baseline controls to exclude any significant differences between CRE-negative and CRE-positive cells in the absence of any other manipulations. We do so in all of our discovery experiments, and have found so far that otherwise unmanipulated CRE-negative and CRE-positive populations have indistinguishable phenotypes.

### Modularity of BEAM enables compatibility across most experimental frameworks

Mosaic analysis is a powerful approach for investigating cell autonomy of gene function, and the modular design of the BEAM system makes it compatible with a wide array of methods for manipulating gene function. Here, we have demonstrated its power using floxed alleles, overexpression, CAS9-mediated knockout and CAS13-mediated knockdown. Moreover, the modular design also makes it possible to modify the proportion of experimental and control cells as needed (*e*.*g*. 90:10, 50:50, 10:90), simply by titrating the dose of *Cre*. With minor modifications, the BEAM plasmid system could also enable spatial and temporal control of gene expression by replacing the *CAG* promoter with tissue-or stage-specific promoters. Alternatively, it also would be possible to restrict analysis to specific cell types of interest using existing *Cre*-driver lines, although this would require a reconfiguration of the BEAM system incorporating a third recombinase to establish an experimental subset of cells within the broader CRE-positive population.

### Applications of BEAM beyond analysis of gene function

Here, we focus on characterizing the BEAM system for investigations of gene function. However, introducing different effector molecules into two distinct cell populations, creates opportunities for many other experimental paradigms. Application of channel rhodopsins (ChRs)(43)(44) or designer receptors exclusively activated by designer drugs (DREADDs)(45)(46), can allow independent stimulation or silencing of specific proportions of neurons in sequence or simultaneously. Anterograde or retrograde synaptic tracing using tetanus toxin c-terminal fragment (TTC)(47)(48) or wheat germ agglutinin (WGA)(49)(50) tagged with distinct epitopes or fluorescent proteins would enable comparative connectomics. Further, in several regenerative paradigms, it might be desirable to instruct production of multiple distinct cell types from a single population of endogenous progenitors, and BEAM could facilitate programming subsets of progenitors toward divergent cell type specification pathways.

In summary, we have developed and validated a highly flexible system for mosaic genetic analysis in mice that will be useful to study a broad range of biological questions in diverse fields of study. The BEAM system is ideally suited to investigation of cell autonomy, as demonstrated here by our analysis of multiple cellular phenotypes controlled by transcription factors during cortical development. However, the BEAM plasmid system should also bring exceptional clarity to subtle biological questions that are difficult to address definitively with existing methods due to experimental variability. An internal control can eliminate or sharply reduce this variability, improving phenotypic identification and reproducibility. Notably, the toolbox of site-specific recombinases has continued to grow in recent years (Anastassiadis et al., 2009; Suzuki et al., 2011; Karimova et al., 2016), raising new and exciting possibilities for combinatorial modulation of gene expression. Therefore, this work lays the foundation for a new generation of increasingly sophisticated recombinase-based tools for mosaic genetic analysis in mice, thus significantly increasing capabilities for discovery and experimental manipulation across a wide range of experimental systems and fields.

## Experimental Procedures

### Construct assembly

All constructs for constitutive mammalian expression were derived by subcloning into the XhoI and StuI sites of CBIG (gift of C. Lois), which contains an abelson murine leukemia virus (A-MuLV) Psi packaging element, a CMV/β-actin promoter, a woodchuck hepatitis virus post-transcriptional regulatory element (WPRE), and an A-MuLV long terminal repeat (LTR). Various configurations of *loxP, lox2272, frt* and *F5* sites were designed *in silico* and produced by GeneArt using gene synthesis, and subcloned into the CBIG backbone. The coding sequences for *EGFP* or *tdTomato* were then subcloned, as appropriate. The coding sequence for *CreM* was obtained from Addgene (Kaczmarczyk and Green JE, 2001; Addgene plasmid #8395). *FlpOM* was generated by introducing the human b-globin intron from *CreM* into a codon optimized *Flp* (Raymond and Soriano, 2007; Addgene plasmid # 13792), interrupting the coding sequence between amino acids 159 and 160. All plasmids generated for this study will be shared on Addgene.

### Mice

All mouse studies were approved by the Harvard University IACUC, and were performed in accordance with institutional and federal guidelines. The date of vaginal plug detection was designated E0.5, and the day of birth as P0. *Satb2*^*fl/fl*^ mice were generated by Grosschedl and colleagues(22)(23), and were generously provided by Susan McConnell. *Couptf1*^*fl/fl*^ mice were generated and generously shared by Studer and colleagues(21)(51). *Rosa26*^*loxP-STOP-loxP-LacZ*^ mice (stock number 003474) were purchased from Jackson Laboratories.

### Immunocytochemistry

Mice were transcardially perfused with 4% paraformaldehyde, and brains were dissected and post-fixed at 4°C overnight in 4% paraformaldehyde. Tissue was sectioned at 50μm on a vibrating microtome (Leica). Non-specific binding was blocked by incubating tissue and antibodies in 8% goat serum/0.3% bovine serum albumin in phosphate-buffered saline. Primary antibodies and dilutions used: mouse anti-SATB2 (1:200, Abcam); chicken anti-β-galactosidase (1:100 Abcam); rabbit anti-GFP (1:500, Invitrogen); rat anti-CTIP2 (1:200, Abcam).

### Imaging

For epifluorescence microscopy, tissue sections were imaged using an Eclipse 90i microscope (Nikon Instruments) with a mounted CCD camera (ANDOR Technology). For confocal microscopy, cells were imaged on an inverted LSM 880 microscope (Zeiss).

### Cell culture and transfection

Human embryonic kidney (HEK) 293T cells were obtained from the ATCC and grown in DMEM (Gibco) containing 10% fetal bovine serum (Seradigm), 5000U/ml penicillin, and 5g/ml streptomycin (Invitrogen). Transfection was performed using Lipofectamine 2000 (Invitrogen) following the protocol provided by the manufacturer.

### In utero electroporation

Surgeries were performed as previously described(28). Briefly, plasmids were microinjected into the lateral ventricle of developing embryos under ultrasound guidance, and electroporated into cortical progenitors using a square wave electroporator (CUY21EDIT, Nepa Gene, Japan) set to deliver five 30V pulses of 50ms, separated by 950ms intervals. For BEAM in utero electroporation experiments all plasmids were used at 1ug/ul each, except for *CAG-Cre*, which was used at 75-125ng/ul (the exact concentration necessary for approximately equal numbers of GFP and RFP labeled cells varies slightly across DNA preps).

### AAV labeling

All virus work was approved by the Harvard Committee on Microbiological Safety, and conducted according to institutional guidelines. We obtained a *pAAV-CAG-EGFP* construct from the MGH Virus Core, which contains the following elements flanked by AAV2 ITRs: a CMV/β-actin promoter, the coding sequence for *EGFP*, the woodchuck hepatitis virus post-transcriptional regulatory element (WPRE), a bovine GH pA signal, and an SV40 pA signal (full sequence available upon request). Mosaic expression *pAAV* constructs were generated by replacing the *EGFP* coding sequence in *pAAV-CAG-EGFP* with the various BEAM modules described above. Constructs were packaged and serotyped with the AAV2/1 capsid protein by the MGH Virus Core. Neurons were labeled by pressure injection of virus under ultrasound guidance at P1, and brains were collected for analysis at P14.

### Fluorescence Activated Cell Sorting

Flow cytometry analysis and FACS purification were performed on a MoFlo Astrios cell sorter (Beckman Coulter) at the Bauer Flow Cytometry Core Facility. Cells were initially gated based on forward and side scatter. Fluorescence gates were set relative to negative controls. Plots in Figure 2C were downsampled to display 1000 fluorescently-labeled cells in each condition, in order to enable direct comparison across samples.

### RNA sequencing

Parietal (somatosensory) cortex was dissected from BEAM electroporated *Couptf1*^*fl/fl*^ brains, acutely dissociated as previously described(52), and FACS purified on a MoFlo Astrios cell sorter at the Harvard Bauer flow cytometry core. Three replicates consisting of 10,000 RFP-or GFP-labeled cells each were obtained from independently electroporated brains. RNA was isolated using RNeasy kit (Qiagen), and libraries were prepared using the SMART-Seq v4 Ultra Low Input RNA kit (TaKaRa). Sequencing was performed on an Illumina NextSeq at the Harvard Bauer core facility. Data were analyzed using Tuxedo tools(53). Gene ontology category enrichment analysis was performed using the PANTHER overrepresentation test(54).

## AUTHOR CONTRIBUTIONS

L.C.G. and J.D.M. conceived the project and designed all experiments. L.C.G. performed all experiments, with M.B.W. contributing to early stages of all experiments.A.P. performed CRISPR-mediated ablation of *Satb2*. S.L. made contributions to cloning and imaging. L.C.G and J.D.M. analyzed and interpreted the data. L.C.G. designed and created figures. L.C.G. and J.D.M. wrote the paper. All authors read and approved the final manuscript.

## ACKNOWLEDGEMENTS

We thank S. Dymecki, S. Ross, R. Grosschedl, S. McConnell, and M. Studer for generous sharing of mice and reagents; P. Davis, E. Gillis-Buck, M. Wettstein and B. Wall for technical assistance; J. Flanagan, S. Dymecki, L. Goodrich, H. Padmanabhan for scientific discussions; and members of the Macklis lab for helpful suggestions. This work was supported by National Institutes of Health grant R21 NS104733 to J.D.M., with additional infrastructure support from NIH DP1 NS106665, NS075672, NS045523, NS104055, and NS049553, Max and Anne Wien Professor of Life Sciences fund, and the Emily and Robert Pearlstein Fund for Nervous System Repair. L.C.G. was partially supported by the Harvard Medical Scientist Training Program, NIH individual predoctoral National Research Service Award NS080343, and the DEARS Foundation (to J.D.M.). M.B.W. was partially supported by NIH individual predoctoral National Research Service Award NS064730 and the DEARS Foundation (to J.D.M.).

## Supplementary Figures

**Figure S1.**
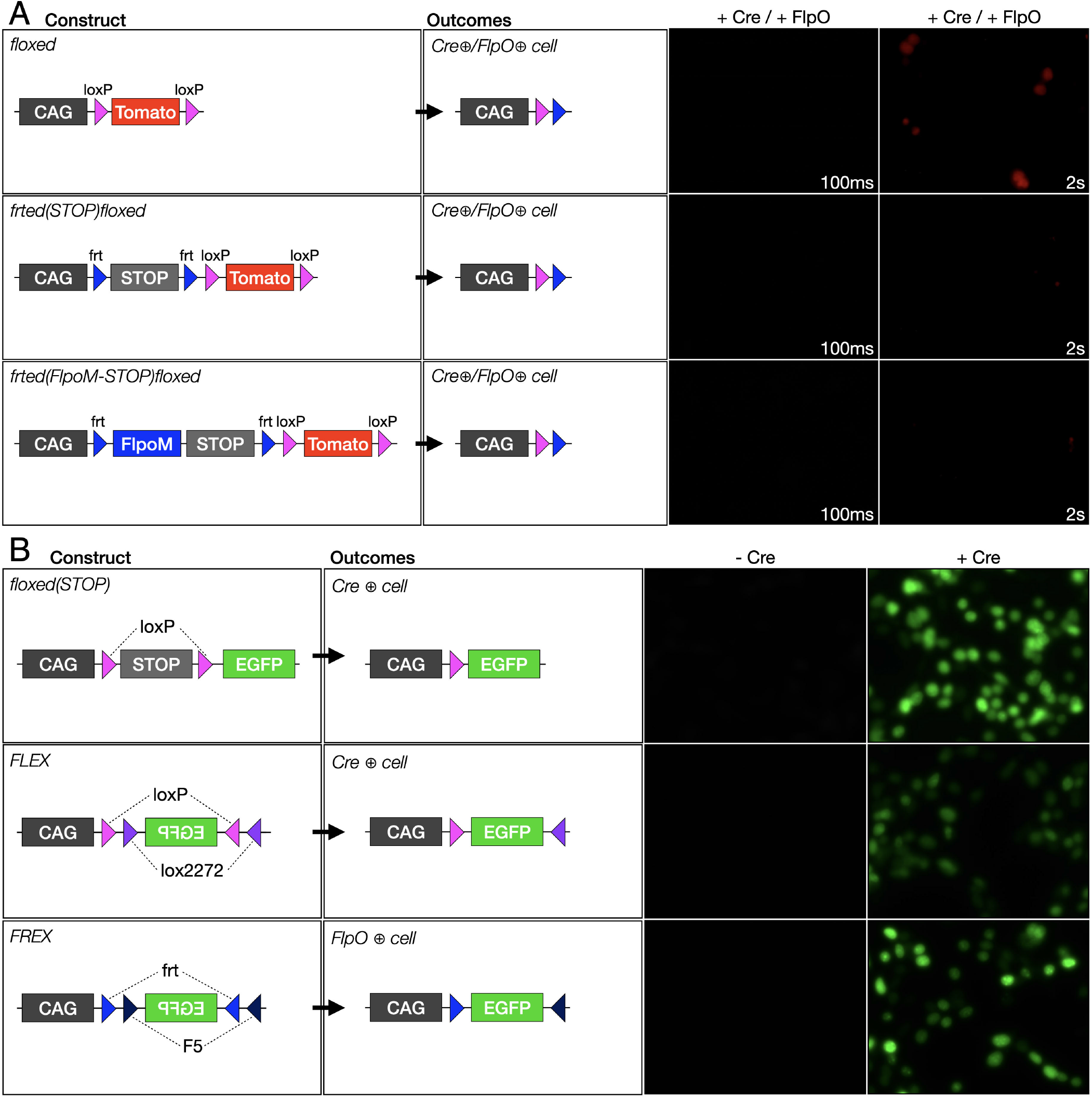
Characterization of recombinase-dependent plasmids for binary gene expression. (A) Reduced RFP expression is observed in CRE-positive cells with Flpo-mediated delay. Co-transfection with a *CAG-Cre* plasmid results in excision of *RFP*, but low levels of RFP are still observed, even in the presence of a high dose of *CAG-Cre*. Delaying *RFP* expression results in direct competition between CRE and FLPO activity, dramatically reducing initial expression of *RFP* in *CAG-Cre*-transfected cells. (B) Strategies for recombinase-dependent gene expression. Excision of a transcriptional *STOP* cassette flanked by direct repeats of a recombination site is highly efficient, but can result in low levels of read-through. Flip excision of inverted incompatible recombinase sites is less efficient, but there is no baseline expression while the gene remains in the antisense orientation.

**Figure S2.**
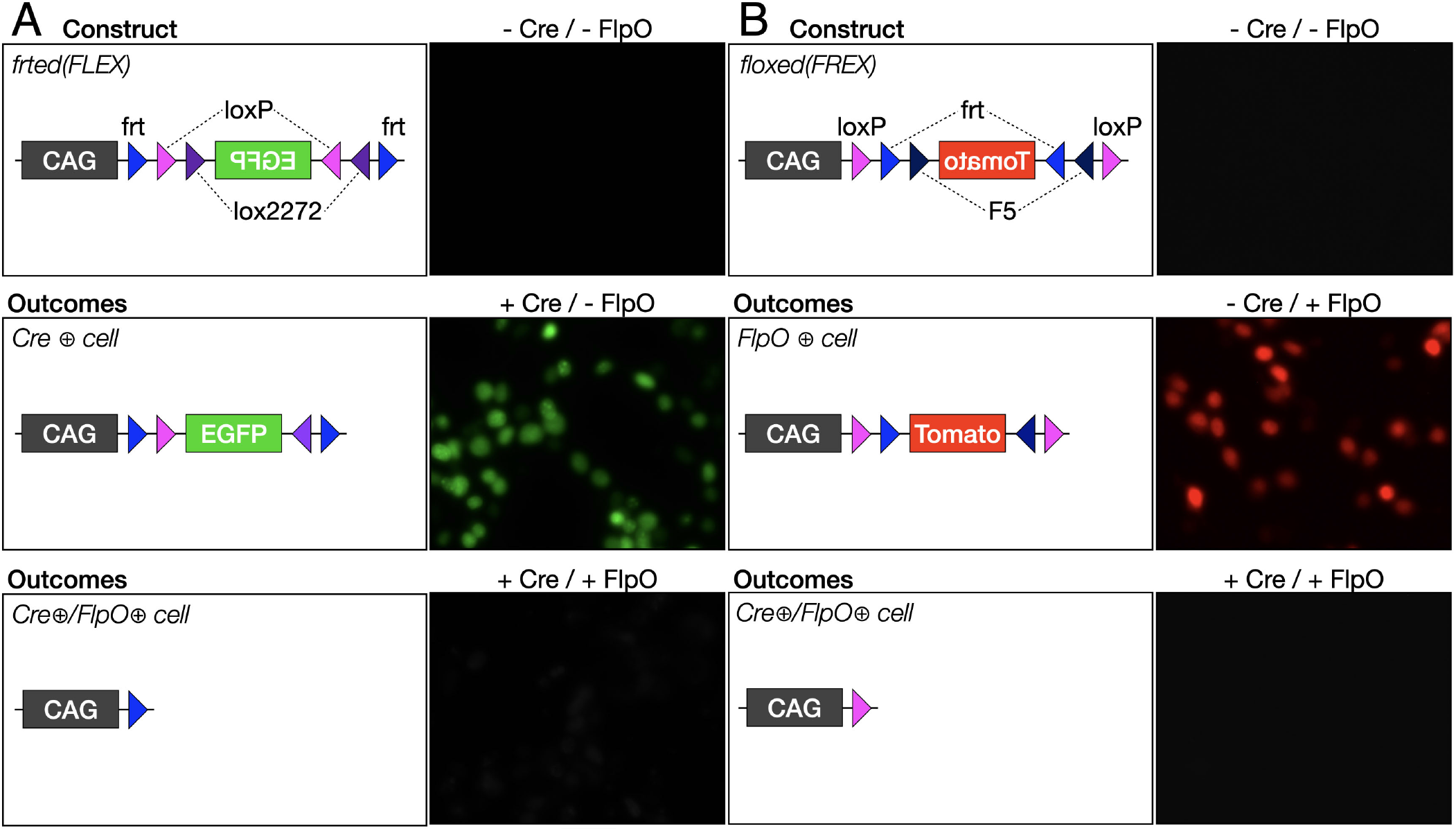
Flip excision constructs for binary gene expression by competitive recombinase activation. (A) In the absence of any recombinase, an inverted *GFP* coding sequence surrounded by *frt* sites and incompatible *lox* sites does not express *GFP* (top). In the presence of CRE, the *GFP* coding sequence is inverted irreversibly, and *GFP* is produced (middle). In the presence of both CRE and FLPO, the *GFP* coding sequence is deleted (bottom). (B) In the absence of CRE or FLPO, an inverted *RFP* coding sequence surrounded by *loxP* sites and incompatible *F* sites does not express *RFP* (top). When CAG-Flpo is added, the *RFP* sequence is inverted and expressed (middle); the presence of both CRE and FLPO leads to deletion of the coding sequence (bottom).

**Figure S3.**
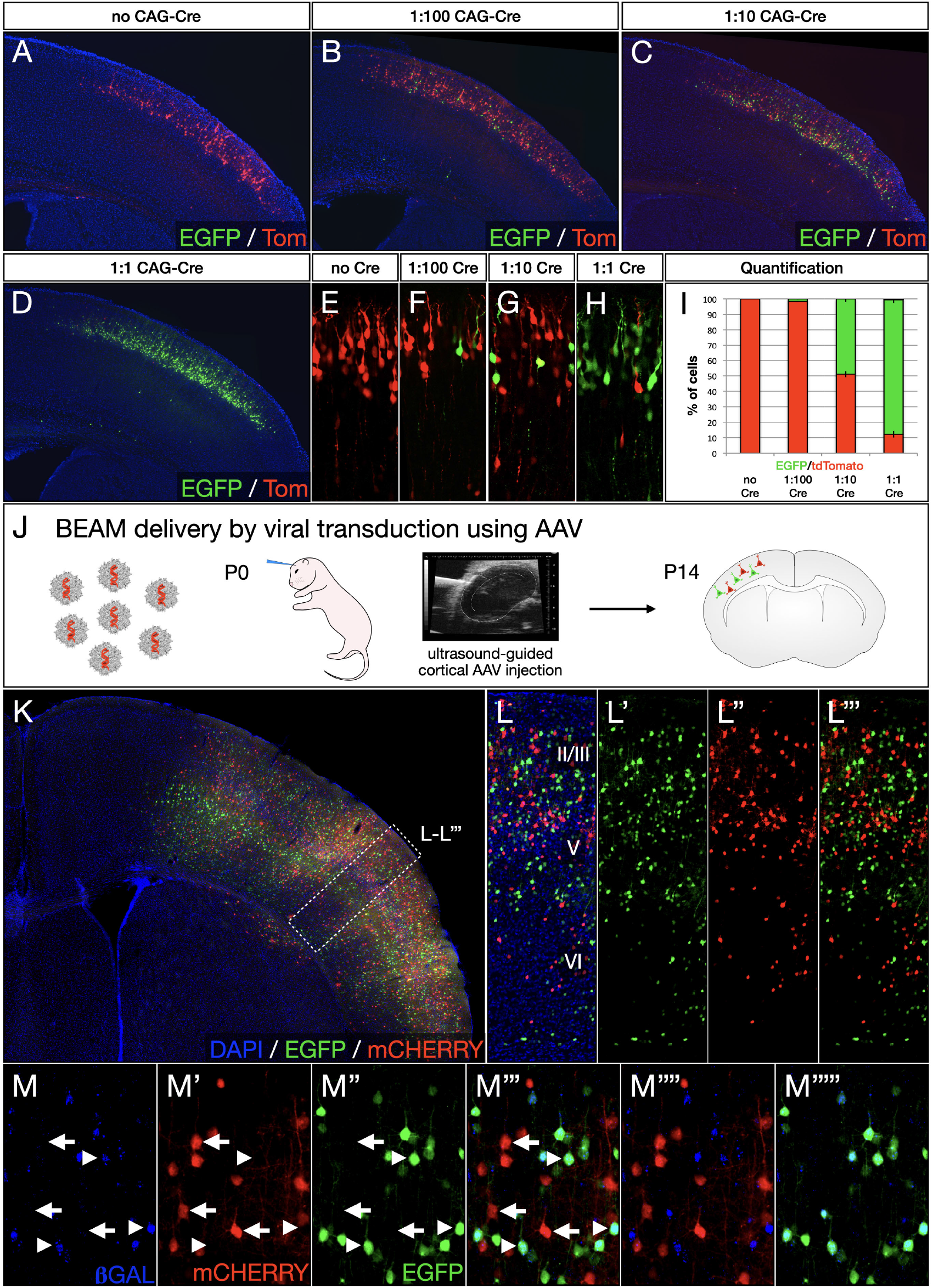
Dose-dependent titration of relative proportions of control and experimental cell populations and AAV-mediated BEAM delivery. (A-I) BEAM plasmids were electroporated at E14.5 with varying concentrations of *CAG-Cre*, from no *CAG-Cre* to 1000 ng/ul *CAG-Cre* (A-D and E-H), resulting in a wide range of ratios of RFP-positive to GFP-positive neurons at P4. Quantification (F). (J-M) Packaging of BEAM into AAV followed by injection into postnatal cortex enables transduction of postmitotic neurons (J) Neurons are labeled across all cortical layers, and distinctly express either GFP or RFP (K-L) Injection into *Rosa26*^*loxP-STOP-loxP-LacZ*^ demonstrates that most GFP-positive neurons are also β-GAL-positive (arrowheads in M-M”””), while most RFP-positive neurons lack β-GAL staining (arrows in M-M”””).

**Figure S4.**
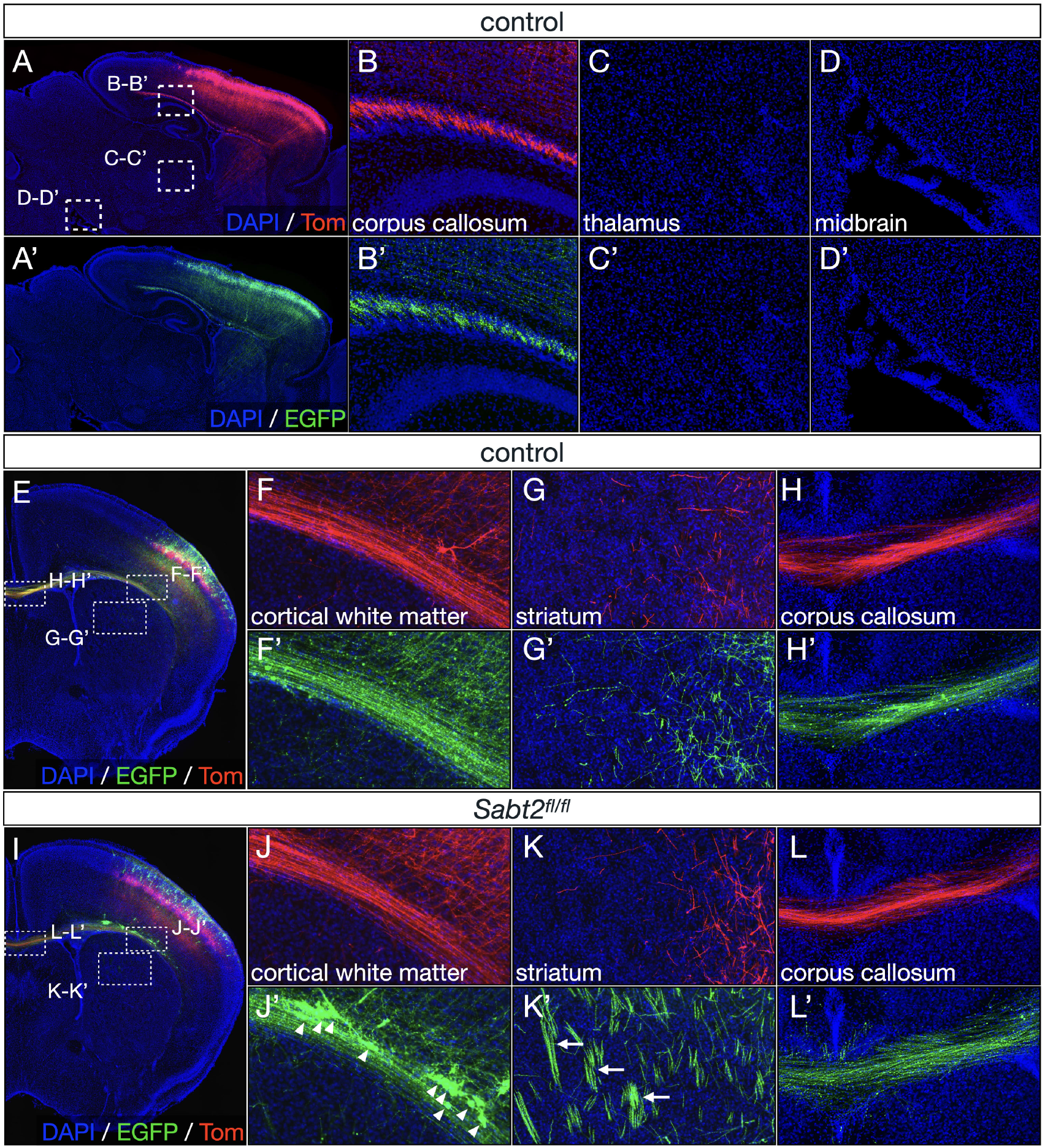
Mosaic deletion of *Satb2* expression by electroporation of BEAM into *Satb2*^*fl/fl*^ mice reveals that the abnormalities of axonal projections are only partially cell-autonomous. (A-H) In control brains electroporated with BEAM plasmids at E14.5 (A-A’), both RFP-positive and GFP-positive superficial-layer neurons send axons through the corpus callosum (B-B’), without any projections to thalamus (C-C’) or midbrain (D-D’). Coronal sections show that both GFP-positive and RFP-positive neurons migrate to superficial layers (F-F’), send only local, branched corticostriatal collaterals in the striatum, without descending fasciculated projections through the internal capsule (G-G’), and project across the corpus callosum (H-H’). (I-J) In *Satb2*^*fl/fl*^ brains (I), RFP-positive neurons continue to migrate to superficial layers (J), send only corticostriatal collaterals (K), and primarily project across the corpus callosum (L). In striking contrast, GFP-positive null neurons frequently remain outside the cortical plate (arrowheads, J’) and many send fasciculated corticofugal axons through the internal capsule (arrows, K’).

**Figure S5.**
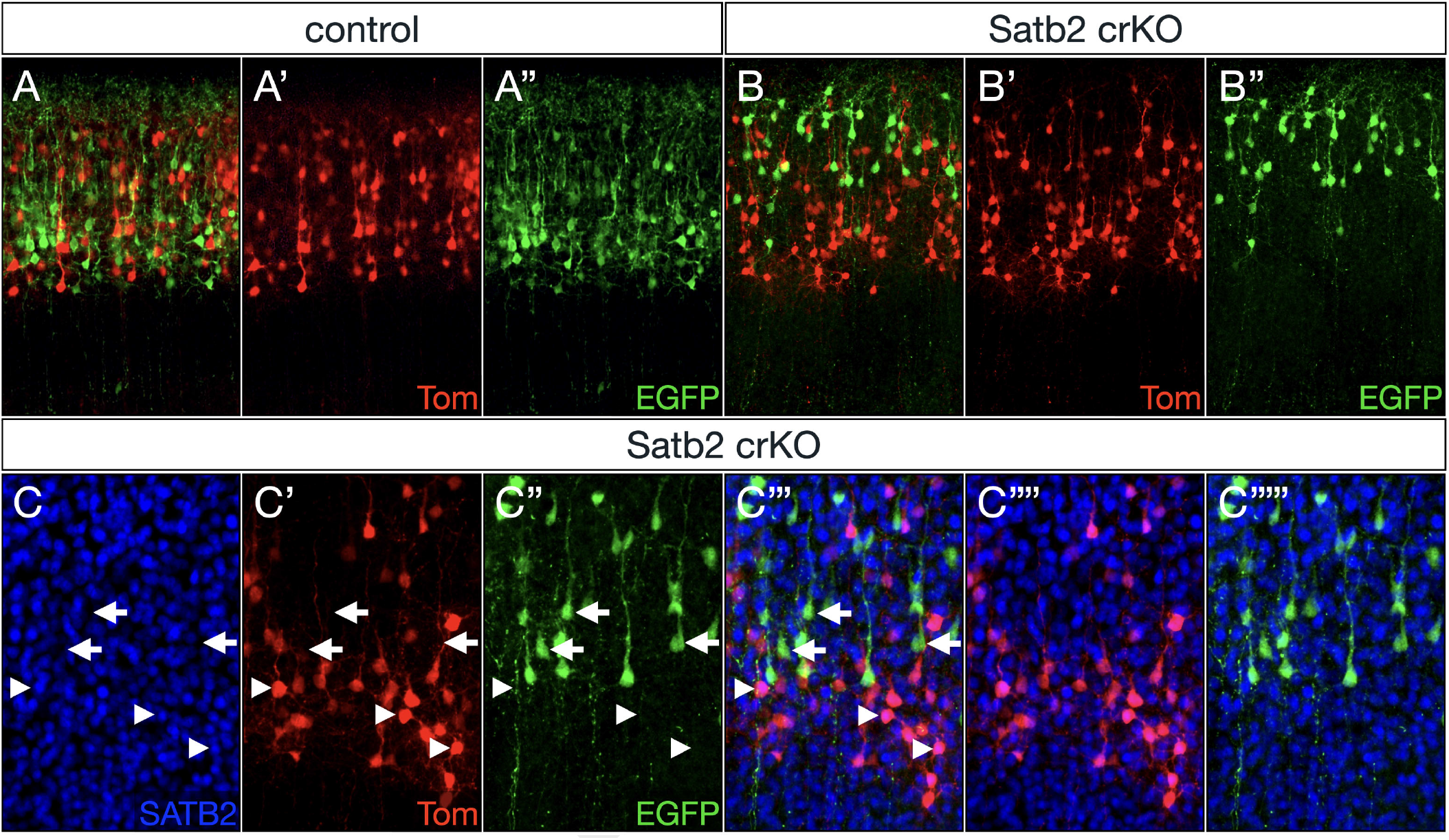
Efficient mosaic gene ablation by CRISPR-Cas9-mediated gene editing using BEAM. (A-B) In control experiments using a guideRNA with no known targets in the mouse genome, both *RFP*- and *GFP*-labeled neurons migrate normally, and reside interspersed in superficial cortical layers (A-A”). In contrast, *Satb2* loss-of-function by CRISPR-Cas9 results in GFP-labeled neurons undergoing a shift to more superficial positions relative to control RFP-labeled neurons. (C) Immunolabeling for SATB2 reveals that almost all GFP-positive cells are SATB2-negative, while almost all control RFP-positive cells are SATB2-positive, indicating that gene ablation is successful and limited to Cre-positive cells that express *Cas9*.

**Figure S6.**
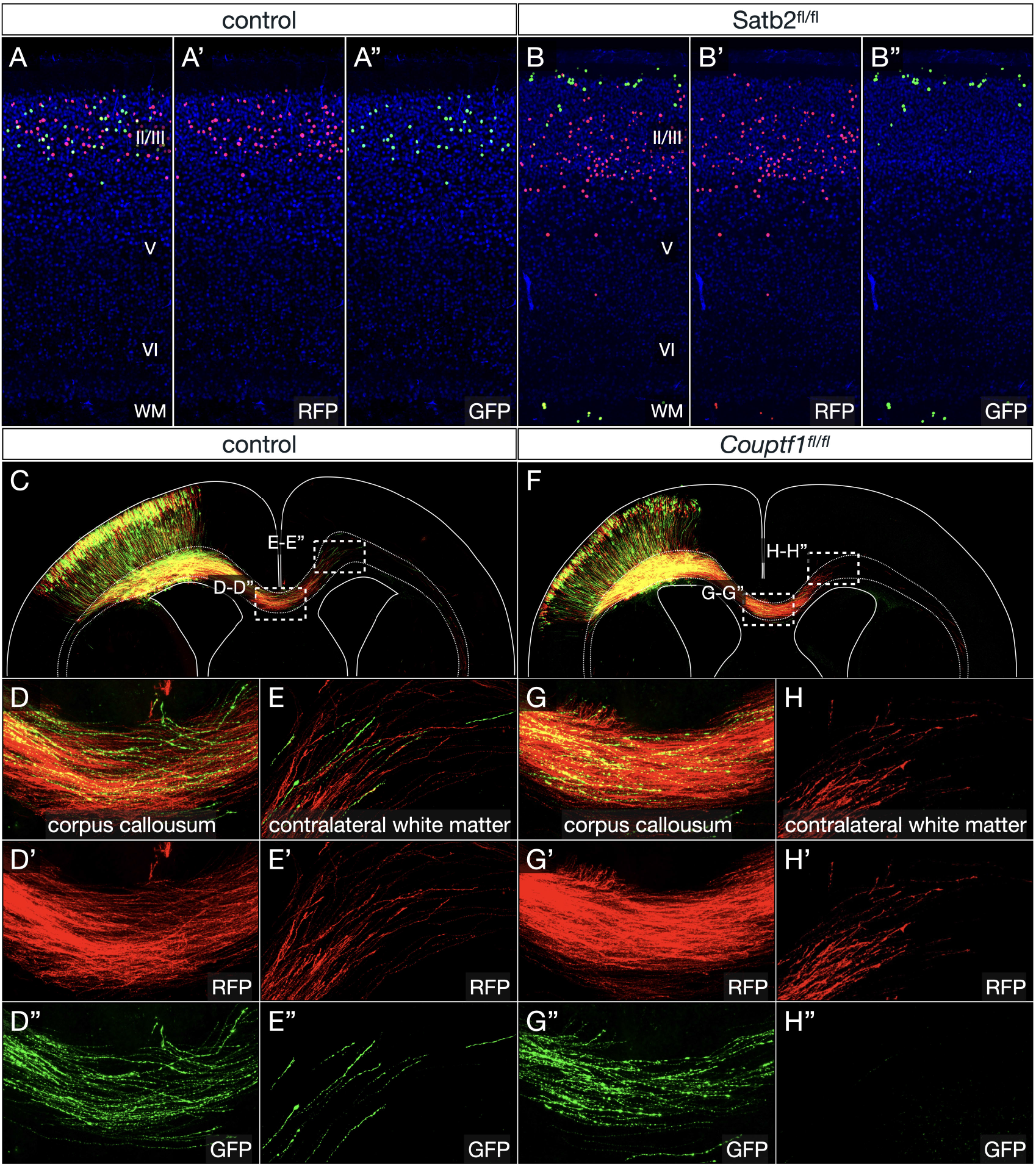
BEAM electroporation into *Satb2*^*fl/fl*^ brains for analysis of final migration patterns and into Couptf1fl/fl brains for analysis of the precise timing of callosal axon extension. (A-B) Mosaic analysis using nuclear BEAM labeling reveals that most *Satb2* conditional null neurons eventually overcome their migrational delay. Analysis of wild-type brains at P7 shows GFP- and RFP-labeled neurons uniformly intermingled across layer II/III (A-A”). In *Satb2*^*fl/fl*^ brains, in contrast, most *Satb2* null GFP-labeled neurons migrate past control RFP-labeled neurons to reside in the most superficial portion of layer II/III, while a smaller subset undergo migrational arrest and remain within the cortical white matter (B-B”). (C-H) BEAM reveals delayed axon extension across the corpus callosum by *Couptf1* conditional null neurons. In wild-type brains, both RFP- and GFP-labeled axons cross the corpus callosum (D-D”) and invade the contralateral cortical white matter (E-E”). In *Couptf1*^*fl/fl*^ brains, control RFP axons behave in the same way, but GFP-labeled axons from *Couptf1* conditional null neurons are delayed reaching the corpus callosum (G-G”), and none have yet reached the contralateral white matter at P0 (H-H”).

**Figure S7.**
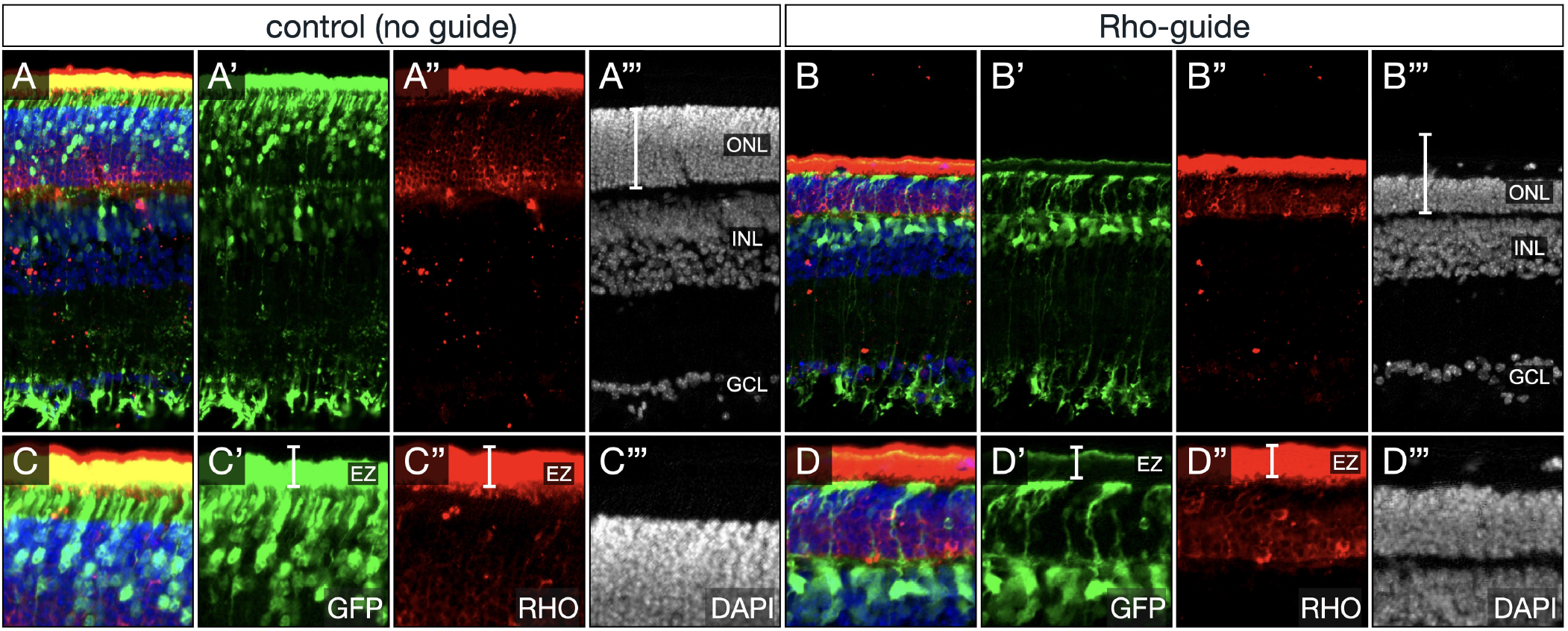
Cas13 enables efficient knockdown of *Rho* in photoreceptors. (A-D) In control brains electroporated at P0 with a *Cas13* expression plasmid but no guide RNA there is normal labeling of cells in the INL and ONL with extensive overlap of GFP and RHO immunostaining at P21 (A-A”‘). In contrast, when expression constructs for guide RNAs targetting *Rho* are also included, very few cells are present in the ONL, which is overall thinner, as seen also by RHO immunostaining (B-B”‘). Note the extensive overlap of photoreceptor outer segments with RHO immunostaning in the elipsoid zone in control experiments (C-C”‘) and the complete absence of overlap in *Rho* experiments (D-D”‘).

**Figure S8.**
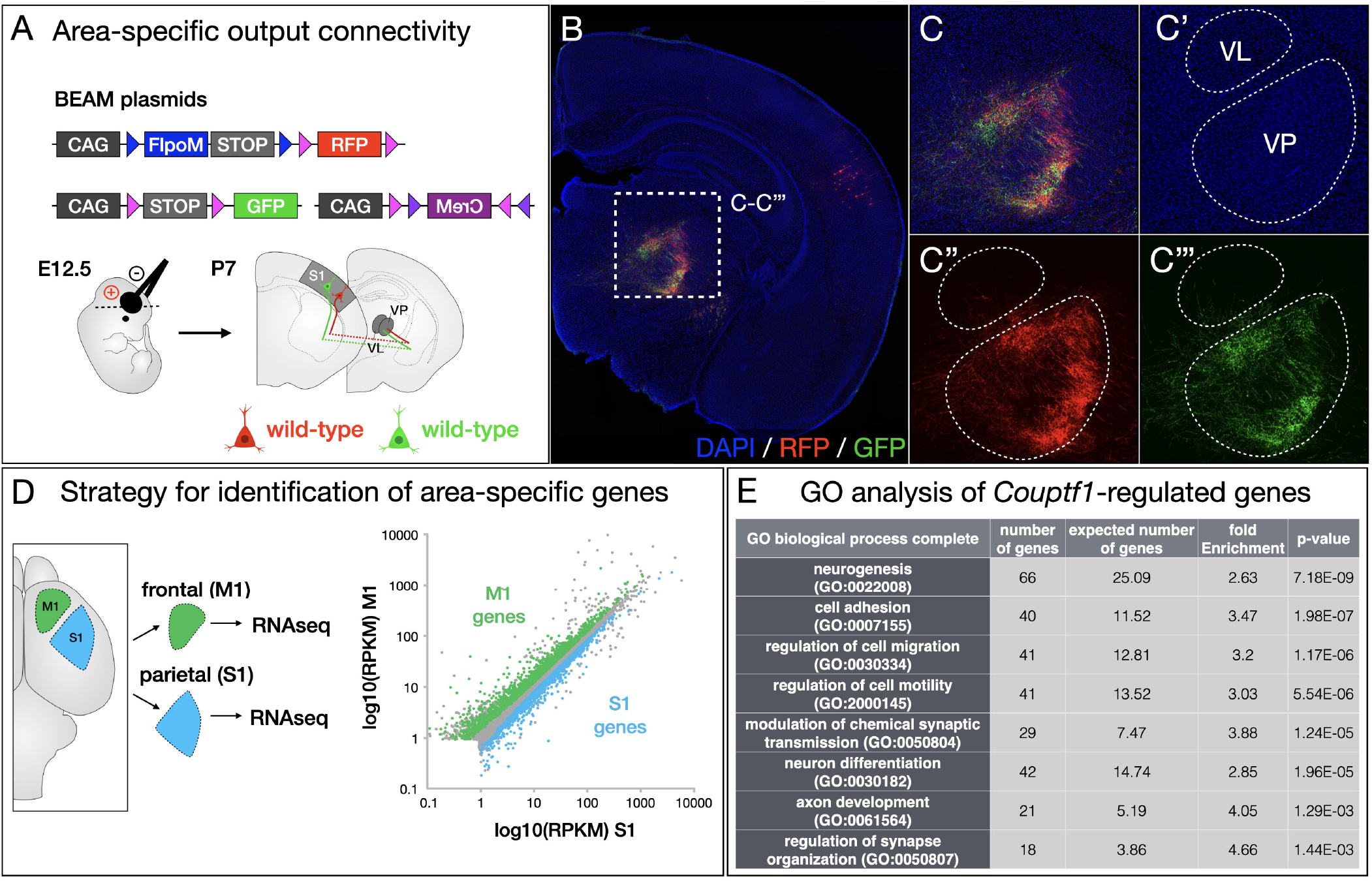
*Couptf1* null cortical columns in sensory cortex aberrantly express genes typical of motor cortex, and establish output connectivity to motor nuclei in the thalamus. (A-C) In control brains electroporated with BEAM plasmids at E12.5 (A), both RFP- and GFP-labeled axons innervate the VP sensory thalamic nucleus with a similar and highly interspersed distribution (C-C”‘). (D) Strategy for RNA-seq identification of genes enriched in motor and sensory cortices in wild-type mice, as previously reported in Greig et al., Neuron 2016. (E) Gene ontology analysis of genes differentially expressed between *Couptf1* wild-type and conditional null neurons indicates that these genes regulate developmental processes that are critical for controlling axonal projection targeting and circuit assembly.

